# Crosslinking of Lipid Nanoparticles Enhances the Delivery Efficiency and Efficacy of mRNA Vaccines

**DOI:** 10.1101/2025.09.03.674081

**Authors:** Xiang Liu, Yining Zhu, Christine Wei, Jinghan Lin, Di Yu, Jiayuan Kong, Fangchi Shao, Jingyao Ma, Tian Xu, Xiaoya Lu, Yunhe Su, Kailei D. Goodier, Leonardo Cheng, Wu Han Toh, Christopher J. Erb, Sixuan Li, Tza-Huei Wang, Hai-Quan Mao

## Abstract

Lipid nanoparticles (LNPs) have enabled the effective delivery of RNA therapeutics and mRNA vaccines. However, their broader applications are limited by the suboptimal stability and endosomal escape efficiency. Here, we present an easy-to-adopt post-assembly crosslinking approach to enhance the structural and functional stability of mRNA LNPs. By leveraging a series of cholesterol derivatives and crosslinking methods, we induce crosslinks of the lipid components following mRNA LNP assembly to form the crosslinked LNPs (cLNPs). We systematically evaluated crosslinking parameters and identified optimal conditions that enhance both the physical stability and transfection efficiency of cLNPs. Our findings demonstrate that cLNPs exhibit improved structural integrity under storage and lyophilization conditions, as well as increased extracellular stability and endosomal escape efficiency, resulting in improved performance of mRNA LNPs both *in vitro* and *in vivo*. This crosslinking strategy represents a critical advance in LNP engineering, enabling more resilient LNPs and broadening the applicability of LNP-based therapies for gene therapy and vaccine delivery. Our work lays the foundation for developing next-generation LNPs with superior stability and delivery efficiency, broadening the impact of RNA therapeutics and vaccines.

## Introduction

Lipid nanoparticles (LNPs) have emerged as a transformative platform for the delivery of nucleic acids, particularly mRNA, marking a new era in vaccine and therapeutic development^1–4^. The success of LNPs was brought into the global spotlight during the COVID-19 pandemic, where mRNA vaccines such as Spikevax (Moderna) and Comirnaty (BioNTech/Pfizer) utilized LNPs to safely deliver mRNA encoding the SARS-CoV-2 spike protein^5–8^. This achievement demonstrated the potential of LNPs in rapid vaccine development and large-scale deployment. Beyond COVID-19 vaccines, LNPs are being intensively researched for a wide range of therapeutic applications including cancer vaccines^9–11^, where they have shown promise in clinical trials targeting various malignancies such as melanoma and solid tumors. By delivering mRNA that encodes for tumor-specific antigens, these LNP-based vaccines aim to stimulate robust immune responses capable of recognizing and attacking cancer cells^12–14^.

Despite their success, the performance of LNPs remains a topic of ongoing research, with efforts focused on improving their delivery efficiency and stability for a broader range of clinical applications^15–17^. Most current strategies aim to enhance LNP performance by engineering new ionizable lipid structures that improve endosomal escape and intracellular delivery^18–20^, optimizing the composition of LNP formulations to achieve better mRNA encapsulation and biodistribution^21–23^, or incorporating additional components for enhanced transfection or tissue targeting ability^24–26^. However, while these approaches have yielded notable advances, maintaining LNP stability and further enhancing endosomal escape remain a significant challenge. In particular, robust structural integrity is the prerequisite for every downstream success: it prevents premature mRNA leakage or degradation during circulation, preserves functional lipids under physiological stress, and reduces compositional drift during large scale manufacturing. A markedly more stable LNP could therefore channel a greater fraction of its payload through the endocytic pathway, amplifying biological potency while simplifying production, storage, and global distribution—especially critical for vaccines and therapeutics destined for resource limited settings.

In this study, we introduce a post-assembly crosslinking strategy for LNP stabilization. This strategy has been previously employed for polymer-based nanoparticles to enhance stability and therapeutic efficacy^27–29^. It is, however, largely unexplored for LNPs due to the delicate balance in identifying the optimal type and degree of crosslinking while permitting sufficient release of mRNA cargo inside the cytosol. Here, we used a series of cholesterol derivatives with different structures and crosslinkers. Following mRNA LNP assembly, the pH is adjusted to 7 to initiate crosslinking, forming crosslinked LNPs (cLNPs). Using a screening workflow, we systematically explored different crosslinking parameters and identified conditions that substantially enhance the stability and transfection efficiency of cLNPs. The physical stability metrics of cLNPs following purification and during the lyophilization process were characterized. Furthermore, the *in vitro* and *in vivo* transfection efficiency of different cLNP formulations was assessed and correlated with the crosslinking degree. We confirmed that this crosslinking approach improves the antigen-specific immune responses of LNP formulations, inducing more effective anti-tumor responses and thereby enhancing the delivery and efficacy of various mRNA-based vaccines and therapies.

## Results

### Construction and screening of crosslinked mRNA LNP formulations

To introduce a crosslinked structure into the LNPs, we designed a crosslinking scheme between diamines and aldehyde or triketone group on cholesterol-derived crosslinker (CDCL), forming cLNPs (**Fig. 1a**). These acid-sensitive cleavable bonds formed under neutral conditions and selectively cleaved at mildly acidic pH (5 – 6). Five CDCL variants (C1–C5) were synthesized, four of which included aldehyde functional groups forming Schiff bases and one of which employed dynamic covalent chemistry involving triketone groups, forming diketone-enamine bonds, as previously reported^30^ (**Fig. 1b-c**). The chemical structures of all five CDCL components were confirmed by ^1^H-NMR spectroscopy (**Supplementary Figs. S1–6**). Building on the FDA-approved SM-102 LNP formulation (molar ratios of SM-102:DSPC:cholesterol:DMG-PEG, 50:10:38.5:1.5) as the starting cLNPs, we incorporated one of the five CDCL variants (C1–C5) and one of the six diamine or polyamine crosslinkers (A1–A6) with varied molar percentages in reference to the total cholesterol. We generated a library of cLNPs by either replacing a fraction of cholesterol with CDCL (minus “-”) or supplementing it with extra CDCL (plus “+”), referred to as “AxCx_n(%)±”. To investigate the effects of crosslinking on stability and performance, we first used firefly luciferase (FLuc) encoding pDNA as a payload surrogate. The stability of these cLNPs was assessed by measuring the released pDNA in response to the treatment with 1.6 µM dextran sulfate (DS). The control groups only contained either CDCL or amino crosslinkers alone. The SM-102 LNP formulation showed a pDNA release of approximately 41.2%, whereas the cLNP formulations consistently exhibited lower cargo release across various CDCL percentages (**Fig. 1d**). Furthermore, cLNPs showed reduced pDNA release as the CDCL percentage increased, indicating enhanced stability with a higher degree of crosslinking. In contrast, LNPs with only CDCL or amino crosslinkers did not yield a reduced release of pDNA. These results confirm effective crosslink with enhanced stability of cLNPs.

**Figure 1.**
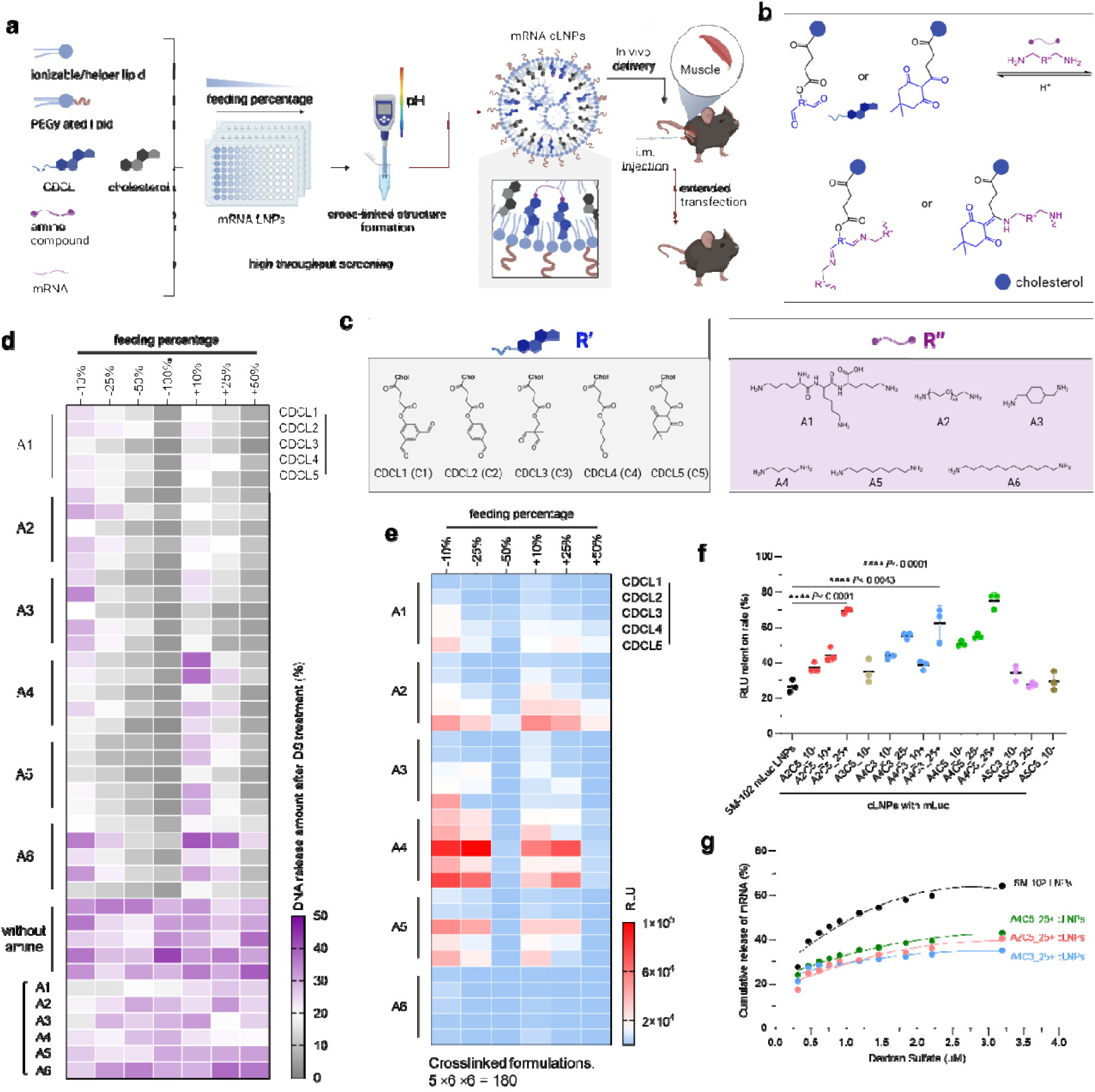
Synthesis, screening, and optimization of crosslinked mRNA LNP formulations. **a.** Schematic of the preparation method of mRNA cLNP formulations. Crosslinkable components such as CDCL and diamine or polyamine crosslinkers, ionizable lipids, helper lipids, cholesterol, and PEGylated lipids, were mixed with the mRNA to form mRNA LNPs. By adjusting the pH, the crosslinking reaction was initiated, resulting in the formation of mRNA cLNP formulations. With further transfection evaluation and stability tests, optimized formulations were identified for further *in vivo* application. **b-c.** Reversible and dynamic, acid-sensitive covalent crosslinks, including diketone-enamine bonds and Schiff base bonds **(b)**, were utilized for the combinatorial design and preparation of cLNP formulations involving structures of five different CDCL and six types of amine crosslinkers **(c)**. **d.** The cLNPs were challenged with 1.6 µM DS solution. The amount of pDNA released from the cLNPs after DS processing is shown in a heat map. **e.** DC2.4 cells were treated with fLuc mRNA cLNPs. The relative luciferase expression after 24 h incubation with fLuc mRNA cLNPs is shown in a heat map. **f.** The retention of transfection efficiency for the 14 top-performing cLNPs were assessed after 24 h of incubation at 37°C. **g.** The cumulative mRNA release from the three cLNPs with optimal stability was evaluated following treatment with different concentrations of DS. Data are from n = 3 (**d**–**g**) biologically independent samples. Data were analyzed using one-way ANOVA for **f**. NS: *P* > 0.05, \**P*L<L0.05, \*\**P*L<L0.01, \*\*\**P*L<L0.001, \*\*\*\**P*L<L0.0001.

To optimize the crosslinking strategy for mRNA LNPs, we first examined the transfection efficiency of the constructed SM-102 cLNP library consisting of 180 formulations on DC2.4 cells, using firefly luciferase encoding mRNA (mLuc). Among the LNPs tested, formulations with CDCL percentages of ±10% and ±25% substantially enhanced mRNA delivery compared to the uncrosslinked SM-102 LNPs. Specifically, compared to the original formulation, A4C3_25-cLNP achieved around a 6.69-fold higher expression level (**Fig. 1e**). Based on this initial screening, 14 mRNA cLNP formulations with the highest transfection efficiencies were selected for further stability tests, where the LNPs were incubated under 37°C for 24 h before dosing. The results showed that mRNA cLNP formulations exhibited substantially improved stability at 37°C with higher transfection efficiency compared to the original formulation. Three mRNA cLNP formulations (A2C5_25+, A4C3_25+, and A4C5_25+ cLNPs), exhibited ∼2.6-fold (*P* < 0.0001), ∼2.3-fold (*P* = 0.0043), and ∼2.8-fold (*P* = 0.0001) higher RLU retention rates than the uncrosslinked SM-102 LNPs, respectively (**Fig. 1f**). The cumulative release of encapsulated mRNA was further analyzed by gradually increasing DS solution concentration, revealing improved stability and a reduced mRNA release profile for these three cLNP formulations compared to the uncrosslinked SM-102 LNP formulation (**Fig. 1g**). Collectively, these results highlight that the three top-performing cLNPs (A2C5_25+, A4C3_25+, and A4C5_25+ cLNPs) have improved both stability and higher transfection efficiency, establishing them as promising candidates for further assessments.

### Optimized cLNP formulations enhance structure stability and mRNA delivery

Following the identification of three top-performing cLNP formulations (A2C5_25+, A4C3_25+, and A4C5_25+ cLNPs), we investigated the physical stability of these cLNPs, particularly during the lyophilization process. Our results showed that compared to original SM-102 LNP formulation, the A2C5_25+ cLNPs retained a consistent hydrodynamic diameter of approximately 252.4 ± 10.2 nm after lyophilization process, with a low polydispersity index (PDI = 0.233 ± 0.018), indicating improved uniformity and structural consistency (**Fig. 2a**). Both the original SM-102 LNP formulation and the A4C3_25+ and A4C5_25+ cLNPs exhibited drastically altered physical properties, with a significant increase in particle size following lyophilization. Further *in vitro* transfection assays revealed that A2C5_25+ cLNPs maintained the same transfection efficiency after lyophilization, whereas the transfection efficiency of the SM-102 LNP formulation decreased by approximately 98.3% following lyophilization. (**Fig. 2b**). Additionally, freeze-thaw cycle testing also demonstrated that only A2C5_25+ cLNPs maintained a PDI below 0.3, even after two or five cycles, preventing nanoparticle aggregation and indicating robust physical stability (**Fig. 2c, d**). The superior stability of the cLNP formulation (A2C5_25+ cLNPs) observed above may result from its unique incorporation of PEG diamine (molecular weight 2000) as the amino crosslinker. This crosslinker yielded the highest degree of protection and structural stability for the cLNPs. These results underscore the resilience of the cLNP formulation to the lyophilization process and freeze-thaw cycles, demonstrating its potential as a stable mRNA LNP formulation for much easier handling and storage.

**Figure 2.**
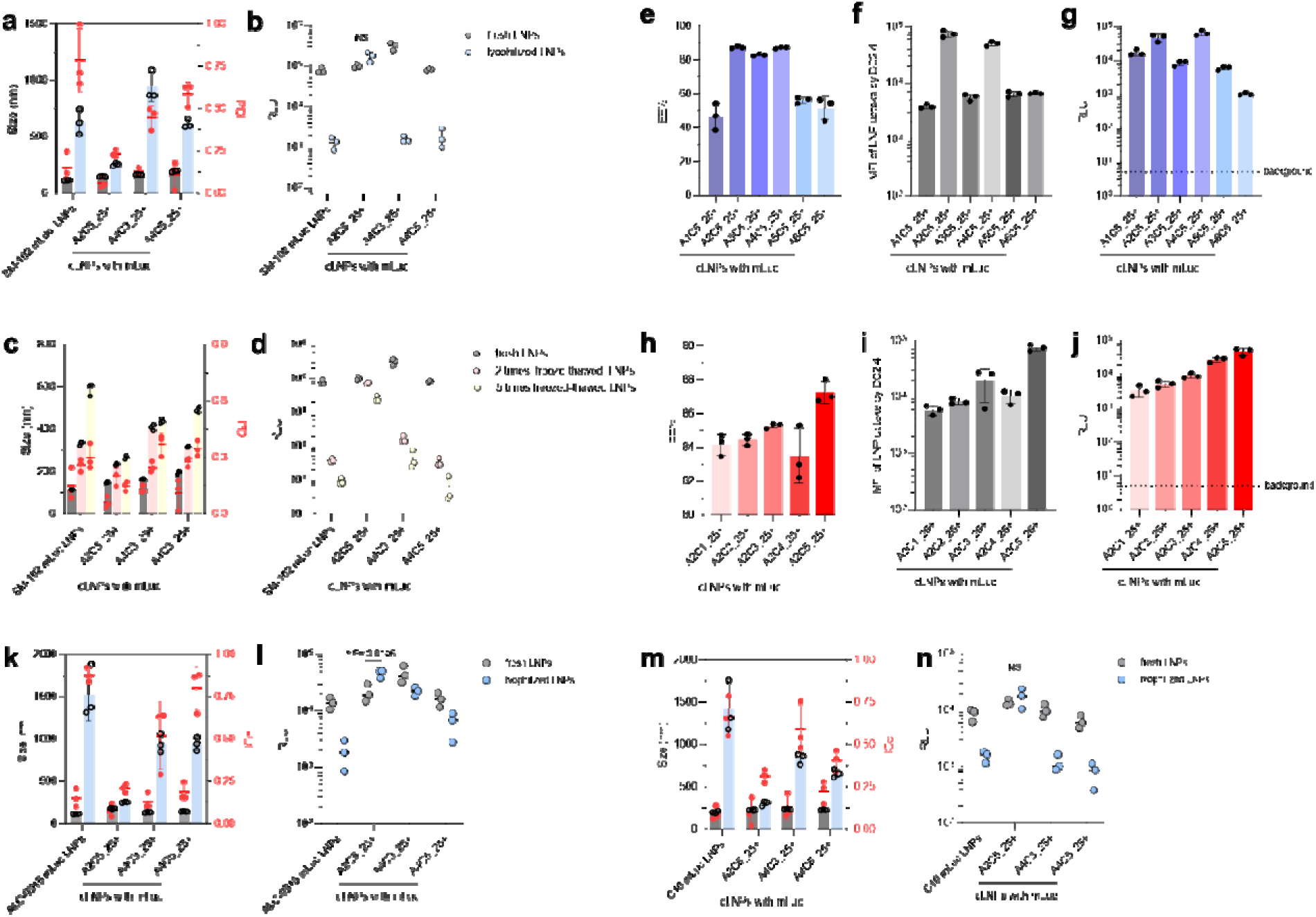
Top-performing cLNP formulations with improved structural stability and transfection efficiency for different LNP formulations. a–b. The particle size distribution **(a)** of the three cLNPs with optimal thermal stability following lyophilization and their transfection efficiency in DC2.4 cells **(b)**. **c–d.** The particle size distribution **(c)** of the three cLNPs with optimal thermal stability following the freeze-thaw cycle and their transfection efficiency in DC2.4 cells **(d)**. **e–g.** The encapsulation efficiency **(e)** of mRNA cLNPs, prepared by fixing CDCL5 and varying 6 amino compounds to create different crosslinking structures, along with the mean fluorescence intensity (MFI) to assess uptake by DC2.4 cells **(f)**, and their transfection efficiency in DC2.4 cells **(g)**. **h–j.** The encapsulation efficiency **(h)** of mRNA cLNPs, prepared by fixing A2 and varying 5 cholesterol derivatives to create different crosslinking structures, along with the mean fluorescence intensity (MFI) to assess uptake by DC2.4 cells **(i)**, and their transfection efficiency in DC2.4 cells **(j)**. **k–l.** The application of three crosslinking ratios to ALC-0315 LNPs formulation, followed by further validation of the particle size distribution **(k)** of the three cLNPs after lyophilization and their transfection efficiency in DC2.4 cells **(l)**. **m–n.** The application of three crosslinking ratios to C10 LNPs formulation, followed by further validation of the particle size distribution **(m)** of the three cLNPs after lyophilization, and their transfection efficiency in DC2.4 cells **(n)**. Data are from n = 3 (**a–n**) biologically independent samples. Data were analyzed using a two-tailed Student’s *t*-test between two groups for **b**, **l**, **n**. NS: *P* > 0.05, \**P*L<L0.05, \*\**P*L<L0.01, \*\*\**P*L<L0.001, \*\*\*\**P*L<L0.0001.

To further explore the structure-function relationship of different cLNPs, we compared the encapsulation efficiency (EE%), cellular uptake, and transfection efficiency of different formulations with specific structures. First, we compared the formulations using the same cholesterol derivative CDCL5 (C5) but with different crosslinkers. Our result showed that, after 24 h of incubation at 37°C, two mRNA cLNP formulations using PEG diamine (A2C5_25+ cLNPs) or 1,4-diaminobutane (A4C5_25+ cLNPs) exhibited the highest EE% values, at 85.4 ± 0.6% and 86.9 ± 0.4%, respectively (**Fig. 2e**). The highest cellular uptake levels were also observed for these two cLNPs (**Fig. 2f**). In addition, these two mLuc cLNP formulations (A2C5_25+, A4C5_25+ cLNPs) also demonstrated the highest levels of transfection efficiency (**Fig. 2g**). These results indicate that, with CDCL5 specifically, these two crosslinkers (PEG diamine or 1,4-diaminobutane) substantially enhanced cLNPs stability, cellular uptake, and transfection efficiency. Second, we fixed the amino crosslinker A2 and varied five types of

CDCL to prepare corresponding cLNP formulations, evaluating the EE% and cellular uptake for each formulation. Among these, the mRNA cLNP formulation using CDCL5 (A2C5_25+ cLNPs) consistently achieved the highest EE% (**Fig. 2h**), cellular uptake (**Fig. 2i**), and transfection efficiency (**Fig. 2j**). These results emphasize the selective interaction between amino crosslinker A2 and crosslinkable CDCL5 and demonstrate that the A2C5 crosslinked structure confers enhanced stability and transfection efficiency, making it a promising configuration for mRNA delivery applications.

Next, we sought to validate the antigen delivery efficiency of the top-performing candidates (A2C5_25+, A4C3_25+, and A4C5_25+ cLNPs) containing mCherry mRNA in bone marrow-derived dendritic cells (BMDCs) using flow cytometry. Compared to the SM-102 LNPs control group, A2C5_25+, A4C3_25+, and A4C5_25+ cLNPs achieved ∼1.72-fold, ∼3.09-fold, and ∼1.99-fold higher transfection efficiency, respectively (**Supplementary Fig. S7a**). To further investigate the immune activation capacity of these cLNPs, we encapsulated ovalbumin-encoding mRNA (mOVA) within each cLNP formulation and assessed BMDC activation and maturation. Compared to SM-102 LNPs, the mOVA cLNPs (A2C5_25+, A4C3_25+, and A4C5_25+ cLNPs) induced elevated CD11c^+^ CD86^+^ SIINFEKL-H-2Kb^+^ expression levels (∼1.17-fold, ∼1.20-fold, and ∼1.17-fold higher, respectively) indicative of enhanced antigen presentation by dendritic cells (**Supplementary Fig. S7b**). These findings suggest the improved transfection efficiency and immune activation potential of the mRNA cLNP formulations, positioning them as strong candidates for mRNA vaccine and immunotherapeutic applications that require robust antigen delivery and immune stimulations.

To confirm the broader applicability and generalizability of our crosslinking strategy, we applied this method to the FDA-approved ALC-0315 LNP formulation and a previously optimized C10 LNP formulation from our previous work^21^. These formulations incorporate distinct ionizable and helper lipids: ALC-0315 LNPs is composed of ALC-0315, DSPC, cholesterol, and DMG-PEG2000 at molar ratios of 46.3:9.4:42.7:1.6, while C10 LNPs contains Dlin-MC3-DMA, DOPE, cholesterol, and DMG-PEG2000 at ratios of 40:40:19.96:0.04. To generate cLNPs, we integrated crosslinkable CDCL and amino crosslinkers at molar amounts aligned with cholesterol content, using the optimized ratios (A2C5_25+, A4C3_25+, and A4C5_25+). The physical stability of these cLNPs post-lyophilization was first assessed. The A2C5_25+ cLNPs preserved the structural quality of ALC-0315 LNPs, maintaining a hydrodynamic diameter of approximately 251.2 ± 10.2 nm and a low polydispersity index (PDI = 0.233 ± 0.018) (**Fig. 2k**). In contrast, the ALC-0315 LNPs without crosslinking exhibited a hydrodynamic diameter of approximately 1523.7 ± 258.7 nm and a remarkably high polydispersity index (PDI = 0.878 ± 0.034), indicating significant particle aggregation and poor uniformity in size distribution. In addition to the differences in the physical properties post lyophilization, ALC-0315 LNPs showed an 86.38% reduction in luciferase expression after lyophilization for uncrosslinked formulation, while the A2C5_25+ cLNPs retained a similar transfection efficiency with a minimal decrease (**Fig. 2l**). Similarly, compared to original C10 LNP formulation, A2C5_25+ cLNPs maintained LNP integrity and transfection efficiency following lyophilization (**Fig. 2m**, **n**). To further evaluate *in vitro* delivery efficacy, we measured mCherry mRNA transfection efficiency in BMDCs using flow cytometry. Both ALC-0315 cLNP and C10 cLNP formulations (A2C5_25+ cLNPs) showed a ∼1.56-fold and ∼1.47-fold increase in transfection efficiency, respectively, over the original formulations (**Supplementary Fig. S8a**). Additionally, the delivery of mOVA mRNA in BMDCs using A2C5_25+ C10 cLNPs induced ∼1.25-fold higher CD11c^+^ CD86^+^ SIINFEKL-H-2Kb^+^ expression levels than the original C10 LNP (**Supplementary Fig. S8b**). Collectively, these results highlight the versatility and generalizability of this crosslinking strategy for different LNP formulations to enhance mRNA delivery stability and transfection efficiency.

### Further physicochemical characterization of the selected mRNA cLNP formulation

Following optimization of the crosslinking conditions, we next sought to characterize the physicochemical properties of the selected crosslinked mRNA LNP formulation (A2C5_25+ cLNPs) (**Fig. 3a**). First, our result revealed that this cLNP formulation exhibited a 31.3% larger average hydrodynamic diameter (∼146.8 nm) compared to that of SM-102 LNPs (∼111.8 nm), along with a ∼2-fold lower PDI. The cLNP formulation also had a nearly neutral surface charge (ζ = −2.88 ± 0.23 mV) and a high mRNA encapsulation efficiency of 92.6%. Representative cryo-transmission electron microscopy (cryo-TEM) images showed that mRNA cLNP formulations retained a spherical shape with a layered shell and an amorphous core, similar to the morphology of original SM-102 LNPs (**Fig. 3b**).

**Figure 3.**
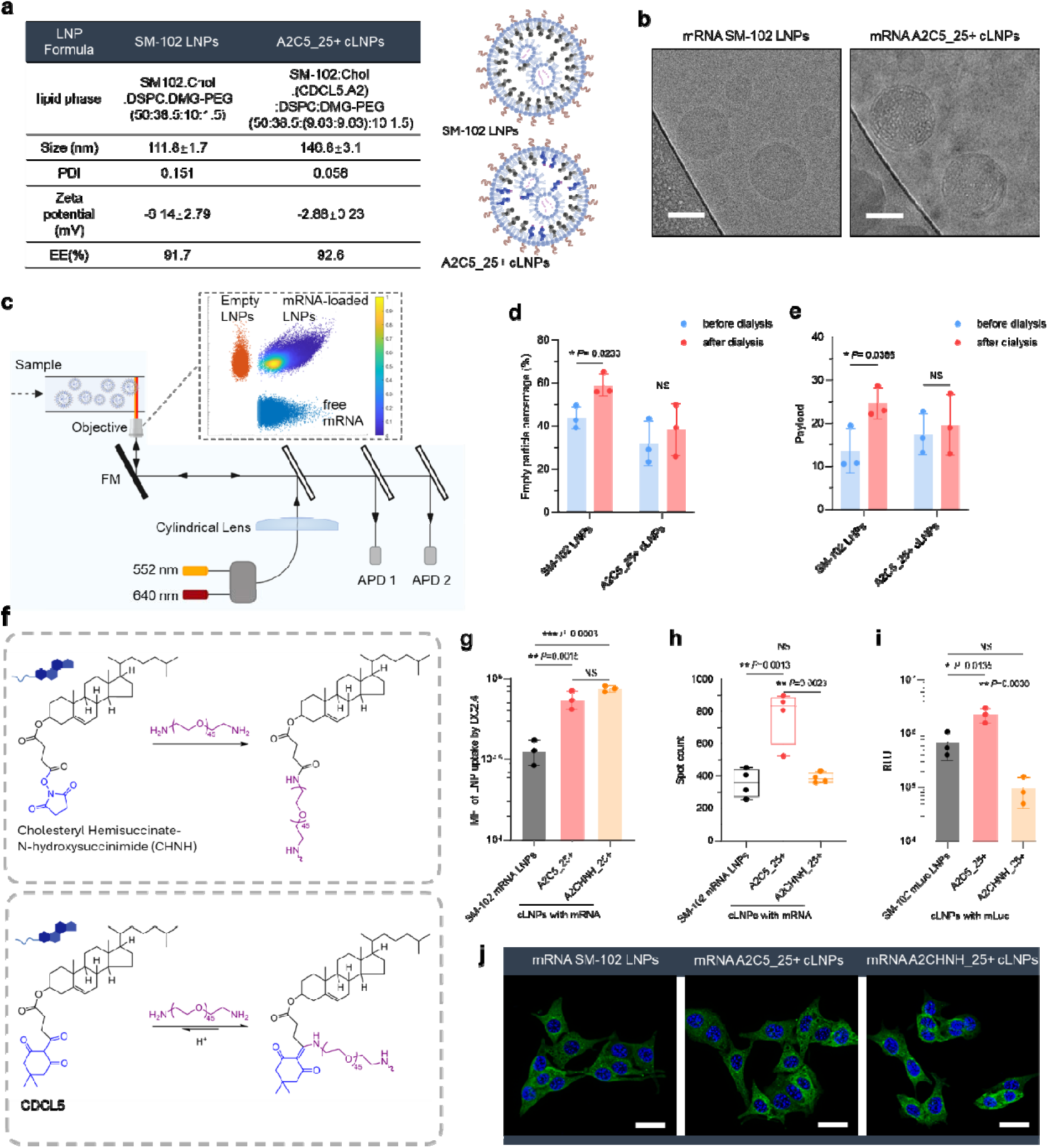
Characterization and enhanced mRNA delivery efficiency of the top-performing crosslinked LNP formulation. **a.** Formulation detail and characterizations of SM-102 LNPs and A2C5_25+ cLNPs. **b.** Representative cryo-TEM images of SM-102 LNP and A2C5_25+ cLNPs. Scale bar: 50 nm. **c.** Schematic illustration of the CICS setup employed for single-particle analysis of LNPs. The LNPs are introduced into a capillary, which traverses a detection light sheet, facilitating fluorescence detection. A cylindrical lens is used to create an observation volume encompassing the entire cross-section of the capillary. As each particle passes through this detection volume, it emits a distinctive fluorescence signal captured by single-photon avalanche photodiodes (APDs) at wavelengths of 552 nm (yellow) and 640 nm (red). The resulting plot of Cy5 and TMR signals—each point representing the fluorescence of a single molecule—enables the discrimination of empty LNPs, mRNA-loaded LNPs, and free mRNA molecules. **d.** The proportion of empty particles in SM-102 LNPs or A2C5_25+ cLNPs before and after dialysis. **d.** Classification of LNPs species into empty particles, mRNA-loaded cLNPs, and free mRNA by plotting TMR signal intensity against Cy5 signal intensity. A representative scatter plot of A2C5_25+ cLNPs following dialysis is provided. **e.** The number-average mRNA copy per SM-102 LNPs or A2C5_25+ cLNPs before and after dialysis. **f.** The structures of the non-cleavable crosslinking component CHNH and the cleavable crosslinking component CDCL5. **g, i.** The MFI **(g)** used to assess the uptake by DC2.4 cells and the transfection efficiency **(i)** in DC2.4 cells for SM-102 LNPs, non-cleavable crosslinked A2CHNH_25+ cLNPs, and cleavable crosslinked A2C5_25+ cLNPs. **h, j.** Quantification (**h**) and representative confocal images (**j**) of GFP-positive spots in B16-Gal8-GFP cells following 4 h of treatment with SM-102 LNPs and cLNPs. Scale bar: 30 μm. Data are from n = 3 (**a**, **d**, **e**, **g**, **i**), n = 4 (**h**) biologically independent samples. For boxplots, the box extends from the 25^th^ to the 75^th^ percentiles, and the line in the middle of the box is plotted at the median. Data were analyzed using a two-tailed Student’s *t*-test between two groups for **d**, **e**, and one-way ANOVA for **g**–**i**. NS: *P* > 0.05, \**P*L<L0.05, \*\**P*L<L0.01, \*\*\**P*L<L0.001, \*\*\*\**P*L<L0.0001.

To assess the impact of crosslinking on compositional stability during dialysis, we utilized a previously developed multi-laser cylindrical illumination confocal spectroscopy (CICS) technique^31^ to examine the mRNA and lipid content in LNP formulations at the single-particle level (**Fig. 3c**). This technique allows for the detection of compositional changes that may occur before and after dialysis. Given that cLNPs form stable crosslinked structures through pH adjustment prior to dialysis, we hypothesized that this increased structural stability would mitigate compositional change. Our results revealed a significant increase in the proportion of empty particles in SM-102 LNPs after dialysis (*P* < 0.05), rising from 43.8 ± 4.3% pre-dialysis to 59.1± 4.3% post-dialysis (**Fig. 3d, Supplementary Fig. S9**). In contrast, the proportion of empty particles in mRNA cLNP formulations remained relatively consistent, with only a subtle increase after dialysis (31.9 ± 8.4% (before) and 38.3 ± 9.8% (after), *P* > 0.05). In the mRNA cLNP formulation, the lower proportion of empty particles compared to SM-102 LNPs can potentially be attributed to an adjusted molar ratio of components, specifically supplementing with extra 25% in CDCL molar ratio relative to the cholesterol content. This adjustment likely hindered the formation of empty particles during the LNPs’ assembly process. Further CICS analysis of mRNA payload distribution revealed that, while SM-102 LNPs exhibited a notable change in the number-averaged mRNA payload among the loaded LNPs (from 13.6 ± 4.2 to 24.9 ± 2.9 following dialysis) that suggest substantial compositional draft, the mRNA cLNP formulations maintained stable mRNA payload levels before and after dialysis process (**Fig. 3e, Supplementary Fig. S10**). These findings, obtained at the single-particle level, highlight the stabilizing effect of the crosslink strategy in minimizing compositional drift, establishing the A2C5_25+ cLNPs as a promising formulation for mRNA delivery.

### Crosslinked mRNA LNPs exhibit enhanced endosomal escape

To investigate the role of acid-sensitive crosslinking in enhancing mRNA delivery, we synthesized a non-cleavable crosslinkable component, cholesteryl hemisuccinate-N-hydroxysuccinimide (CHNH), incorporating active ester functionalities (**Fig. 3f, Supplementary Fig. S11**). CHNH was then reacted with an A2 amino crosslinker to establish robust amide bonds, forming non-cleavable A2CHNH_25+ cLNPs. Cellular endocytosis assays revealed significant increases in the internalization of both A2C5_25+ cLNPs and A2CHNH_25+ cLNPs, with uptake enhancements of ∼2.1-fold and ∼2.4-fold, respectively, compared to the uncrosslinked SM-102 LNPs (**Fig. 3g**). These results underscore the role of crosslinking structures in facilitating cellular uptake.

Considering the critical role of endosomal escape in the mRNA delivery process^32, 33^, we next evaluated the impact of acid-sensitive crosslinked structures on LNP-mediated endosomal escape. We utilized the B16-Gal8-GFP cell line, which recruits Gal8-GFP to sites of endosomal membrane disruption, thereby providing a means to visualize and quantify successful endosomal escape^34^. We observed a notable increase in endosomal escape with the acid-sensitive crosslinked mRNA LNPs compared to the uncrosslinked SM-102 LNPs. Specifically, the acid-sensitive A2C5_25+ cLNPs facilitated ∼2.2-fold greater endosomal escape than SM-102 LNPs (**Fig. 3h**, **j**). The enhanced endosomal escape observed with acid-sensitive A2C5_25+ cLNPs is likely due to their pH-responsive crosslinked structure, which destabilizes under acidic endosomal conditions, facilitating efficient mRNA release into the cytoplasm. In contrast, the non-cleavable A2CHNH_25+ cLNPs, while showing improvements in intracellular uptake, did not enhance endosomal escape. This highlights the essential role of cleavable crosslinking structures in promoting mRNA release from the endosomal compartment.

Further, *in vitro* transfection studies using DC2.4 cells demonstrated that acid-sensitive A2C5_25+ cLNPs significantly enhanced gene expression levels, achieving a ∼3.3-fold increase relative to SM-102 LNPs (**Fig. 3i**). However, the non-cleavable A2CHNH_25+ cLNPs seemed to reduce the transfection efficiency. These findings demonstrate that the cleavable crosslinked network within the LNPs promotes cellular uptake, and increases endosomal escape capability, leading to a higher transfection efficiency.

### Crosslinked mRNA LNPs for sustained and higher *in vivo* mRNA expression

To investigate the *in vivo* mRNA transfection efficiency and expression duration of the mRNA cLNP delivery system, we first selected the A2C5 crosslinking structure based on its enhanced *in vitro* stability and improved endosomal escape. C57BL/6 mice were injected intramuscularly (*i.m.*) with either uncrosslinked SM-102 LNPs or A2C5 cLNPs loaded with mLuc, and luciferase expression was monitored daily using *in vivo* bioluminescence imaging (IVIS). Our result revealed that the mRNA cLNP formulations (A2C5_10+ and A2C5_25+ cLNPs) exhibited 30-fold and 71-fold higher luciferase expression, respectively, than the uncrosslinked SM-102 mRNA LNPs (**Fig. 4a**, **b**). In addition to the more potent transfection efficiency, a more sustained luciferase expression was observed. For A2C5_10+ and A2C5_25+ cLNPs, transgene expression persisted for 15 and 19 days, respectively, whereas the SM-102 LNPs exhibited luciferase activity for only 7 days (**Fig. 4c**). These findings suggest that the optimized cLNP formulations not only achieved higher transfection efficiencies but also extended the duration of gene expression.

**Figure 4.**
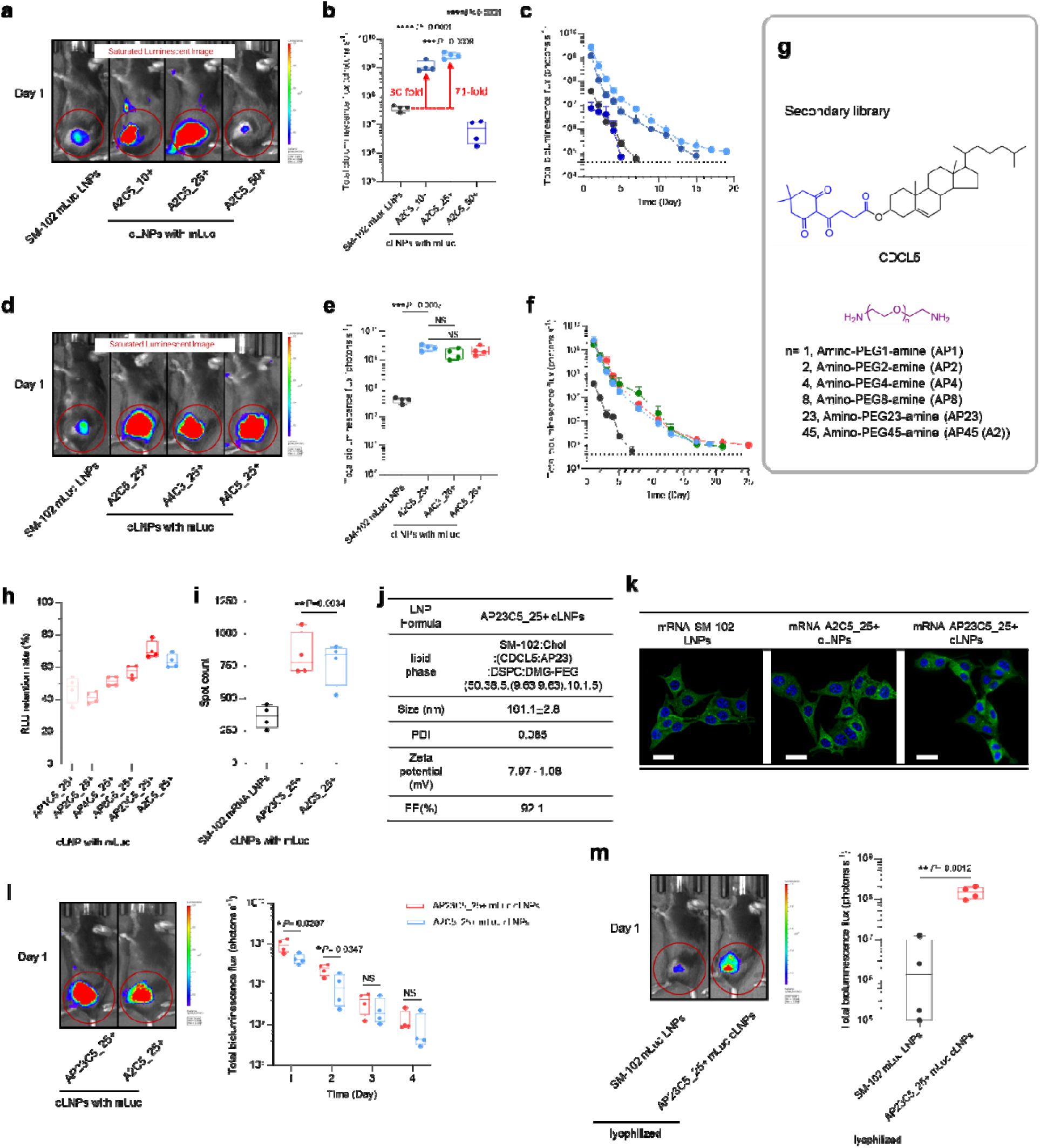
Crosslinked LNP-mediated mRNA delivery *in vivo*. **a.** SM-102 LNPs and three different feed percentages of A2C5 cLNP formulations were *i.m.* injected into mice, and luciferase expression was imaged daily post-injection using IVIS (n = 4 biologically independent samples). A representative bioluminescence image from Day 1 is presented. **b.** The total luminescent flux at the injection site was quantified on Day 1 post-injection. **c.** Quantification of total bioluminescent flux at the injection site in mice measured at different time points post-injection. **d.** SM-102 LNPs and three different cLNP formulations were i.m. injected into mice, and luciferase expression was imaged daily post-injection using IVIS (n = 4 biologically independent samples). A representative bioluminescence image from Day 1 is presented. **e.** The total luminescent flux at the injection site was quantified at Day 1 post-injection. **f.** Quantification of total bioluminescent flux at the injection site in mice measured at different time points post-injection. **g.** The structures of CDCL5 and PEG diamine with different molecular weights. **h.** The retention rates of RLU for the six cLNPs with different fractions of PEG diamine were assessed after 24 h of incubation at 37 °C. **i, k.** Quantification **(i)** and representative confocal images **(k)** of GFP-positive spots in B16-Gal8-GFP cells following 4 h of treatment with AP23C5_25+ cLNPs. **j.** Formulation detail and characterizations of AP23C5_25+ cLNPs. **l.** A2C5_25+ and AP23C5_25+ cLNP formulations were *i.m.* injected into mice, with luciferase expression at the injection site imaged and quantified daily for total luminescent flux over the first four days post-injection using the IVIS (n = 4 biologically independent samples). Representative bioluminescence images from Day 1 is presented. **m.** Lyophilized SM-102 LNP and AP23C5_25+ cLNP formulations were *i.m.* injected into mice, with luciferase expression at the injection site imaged and quantified for total luminescent flux on Day 1 post-injection using the IVIS (n = 4 biologically independent samples). A representative bioluminescence image from day 1 is presented. Data represent meanL±Ls.e.m. with n = 4 (**a**–**f, h**–**i, k**–**m**), n = 3 (**j**) biologically independent samples. For boxplots, the box extends from the 25th to the 75th percentiles, and the line in the middle of the box is plotted at the median. Data were analyzed using one-way ANOVA for **b**, **e**, **h**, **i**, and two-tailed Student’s *t*-test between two groups for **l** and **m**. NS: *P* > 0.05, \**P*L<L0.05, \*\**P*L<L0.01, \*\*\**P*L<L0.001, \*\*\*\**P*L<L0.0001.

Next, we further investigated the effect of different crosslinking structures on mRNA delivery efficiency. Mice were *i.m.* injected with mLuc cLNP formulations consisting of various crosslinking structures (A2C5_25+, A4C3_25+, or A4C5_25+ cLNPs), and luciferase expression was monitored daily. Day 1 results showed that A2C5_25+, A4C3_25+, and A4C5_25+ cLNPs all showed a marked elevated transfection potency than the original SM-102 formulation (around 71-fold, 44-fold, and 56-fold higher, respectively) (**Fig. 4d**, **e**). In addition, in contrast to the uncrosslinked SM-102 LNPs, sustained luciferase expression was observed for A2C5_25+, A4C3_25+, and A4C5_25+ cLNPs, lasting up to 19, 21, and 25 days, respectively (**Fig. 4f**).

### Refining crosslinked LNPs for optimal *in vivo* mRNA delivery

To further optimize the A2C5 crosslinking structure for enhanced *in vivo* mRNA delivery, we modified the PEG diamine component by varying its molecular weight, *i.e*., the ethylene oxide repeat units (n = 1, 2, 4, 8, 23, 45), while keeping the crosslinkable CDCL5 constant. This resulted in six crosslinked mRNA LNP formulations designated as AP1C5_25+, AP2C5_25+, AP4C5_25+, AP8C5_25+, AP23C5_25+, and A2C5_25+ cLNPs (**Fig. 4g**). These variations allowed us to systematically investigate the effect of molecular weight changes on cLNP performance. Each formulation, loaded with mLuc, was incubated at 37 °C for 24 h before dosing, and the RLU retention was then calculated. Among the tested crosslinked formulations, AP23C5_25+ cLNPs exhibited the highest RLU retention rate, suggesting that the molecular weight of the PEG diamine substantially influenced the stability and transfection efficiency of the cLNPs (**Fig. 4h**). In addition, quantification of GFP-positive spots in cells treated with AP23C5_25+ cLNPs showed a ∼2.3-fold increase in endosomal escape compared to SM-102 LNPs (**Fig. 4i, k**). The physicochemical properties of AP23C5_25+ cLNPs are shown in **Fig. 4j**. AP23C5_25+ cLNPs exhibited a hydrodynamic diameter of ∼161.1 nm with a PDI of 0.085, indicating a narrow size distribution. Additionally, they displayed a negative surface charge (ζ = −7.97 ± 1.08 mV) and a high mRNA encapsulation efficiency of 92.1%. These results highlight the important role of molecular weight in optimizing intracellular delivery. Among the formulations tested, the AP23C5_25+ cLNPs demonstrated superior stability and intracellular delivery potential due to the more optimal crosslinker structure.

To determine the effect of this structural refinement on *in vivo* mRNA transfection efficiency, we compared the *in vivo* transfection efficiency of AP23C5_25+ cLNPs and the previously optimized A2C5_25+ cLNPs. C57BL/6 mice were *i.m.* injected with mLuc-loaded crosslinked AP23C5_25+ or A2C5_25+ cLNPs, and luciferase expression was systematically monitored daily over the initial four days post-injection to capture early transfection dynamics. Our result showed that, on Days 1 and 2, AP23C5_25+ cLNPs induced ∼2.2-fold (*P* = 0.0297) and ∼2.5-fold (*P* = 0.0347) higher gene expression levels than A2C5_25+ cLNPs, respectively (**Fig. 4l**), suggesting an improvement in mRNA delivery efficiency with the refined AP23C5_25+ crosslinking structure. We further explored *in vivo* mRNA delivery efficacy of lyophilized crosslinked AP23C5_25+ cLNPs and found that the mLuc-loaded crosslinked AP23C5_25+ cLNPs exhibited a ∼38-fold enhancement in gene expression levels post-lyophilization compared to their corresponding uncrosslinked SM-102 LNP formulations (**Fig. 4m**). In addition, the histological staining of major organs such as heart, liver, spleen, lung, and kidney showed that AP23C5_25+ cLNPs did not induce obvious inflammatory cell infiltration and pathological changes (**Supplementary Fig. S12**), indicating that the crosslinkable compounds have good biocompatibility *in vivo*.

Further, we applied the optimized AP23C5_25+ ratios to the ALC-0315 and C10 LNP formulation as detailed in the molar proportions outlined in **Supplementary Table 1**. C57BL/6 mice were *i.m.* injected with mLuc-loaded crosslinked LNPs and original LNPs, and luciferase expression was monitored on the first day post-injection. Our results indicate that gene expression levels induced by AP23C5_25+ cLNPs were ∼5-fold higher than those induced by the original ALC-0315 LNPs (*P* = 0.0197) (**Supplementary Fig. S13**). Similarly, gene expression levels induced by AP23C5_25+ crosslinked LNPs were ∼9-fold higher than those induced by the original C10 LNPs (*P* < 0.0001) (**Supplementary Fig. S14**). These findings demonstrate that structural optimization in the mRNA cLNP formulations enhances mRNA transfection potency and duration *in vivo*, and the AP23C5_25+ cLNP formulation is shown to be a more efficient delivery system. Consequently, crosslinked AP23C5_25+ cLNPs were selected for further immune assessments.

### Enhanced antigen-specific immune response induced by mRNA cLNPs

To evaluate the vaccine potentcy of the mOVA-loaded AP23C5_25+ cLNPs *in vivo*, we vaccinated mice on Days 0, 7, and 14, using mOVA-loaded uncrosslinked SM-102 LNPs as a control (**Fig. 5a)**. On Day 21 post-vaccination, spleens were collected for T-cell activation assessment. As **Fig. 5b**, **Supplementary** Figs. S15, 16, and **Fig. 5e** showed, crosslinked mRNA LNP formulations led to a substantially elevated increase (∼2.0-fold) in the frequency and number of OVA^+^ CD8^+^ T cells in the spleen, compared to the original SM-102 LNP, indicating an enhanced antigen-specific CD8^+^ T cell response by the selected cLNP formulation. To further assess T cell functionality, lymphocytes isolated from the spleens were subjected to *in vitro* antigen restimulation for intracellular cytokine staining, and approximately 2.3-fold and 3.1-fold increases were observed in the frequencies of CD3^+^ CD8^+^ IFN-γ^+^ (**Fig. 5c**, **Supplementary Figs**. **S17**, **18**) and CD3^+^ CD8^+^ TNF-α^+^ (**Fig. 5d, Supplementary Fig**. **S19**) cells, respectively, compared to SM-102 LNP treated group. After restimulation with antigen, there was a ∼2.3-fold increase in the production of the pro-inflammatory cytokine IFN-γ relative to SM-102 LNPs (**Fig. 5f**). This enhancement further signifies that the cLNP formulation successfully elicited a potent antigen-specific cytolytic T cell-mediated responses. Humoral immune responses were further examined for the selected AP23C5_25+ cLNPs. Compared to SM-102 LNPs, the AP23C5_25+ mRNA cLNPs elicited higher OVA-specific IgG titers, showing a ∼94-fold increase in total IgG (**Fig. 5g**) levels as well as ∼13-fold and ∼4-fold increases in IgG1 (**Fig. 5h**) and IgG2C (**Fig. 5i**) subclass titers, respectively. These results indicate that the mRNA cLNP formulation successfully induced a more potent antigen-specific cellular and humoral immune response than the original SM-102 LNP formulation.

**Figure 5.**
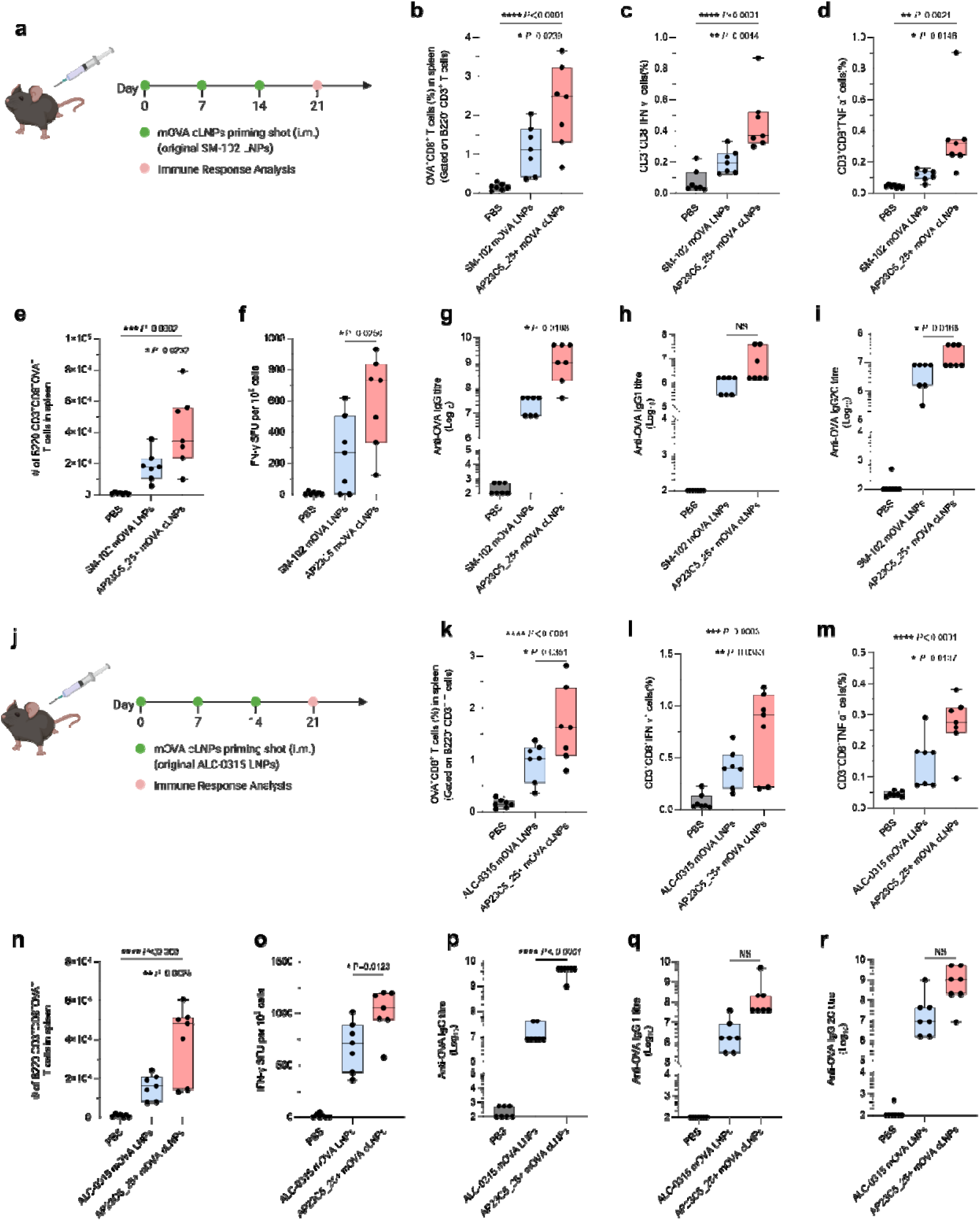
*In vivo* assessment of crosslinked LNP formulations for enhanced mRNA vaccine immunity. a–d. C57BL/6 mice were administered with PBS, SM-102 LNPs, or crosslinked AP23C5_25+ cLNPs (based on SM-102 LNPs) loaded with mOVA via *i.m.* injection (10Lμg mOVA per injection). Mice were sacrificed 7 days after the final injection, and their splenocytes were isolated for analysis **(a)**. The percentages of OVA-specific CD8 T cells (B220^-^CD3^+^CD8^+^OVA^+^ cells) **(b)**. Splenocytes were restimulated in vitro with OVA and SIINFEKL peptide (100LμgLml^−1^ OVA and 2LμgLml^−1^ SIINFEKL) for 6Lh and assessed via flow cytometry and intracellular cytokine staining to determine the percentages of CD3^+^CD8^+^IFN-γ^+^ **(c)** and CD3^+^CD8^+^TNF-α^+^ **(d)**. **e.** The number of B220^-^CD3^+^CD8^+^ OVA-specific T cells in the spleen was analyzed through flow cytometry. **f.** Frequency of IFN-γ-producing cells in spot-forming unit (SFU) among restimulated splenocytes, assessed via ELISpot. **g–i.** Titers of OVA-specific IgG **(g)**, IgG1 **(h)**, and IgG2c **(i)** antibodies in blood serum on Day 21, determined by ELISA. **j**–**m.** C57BL/6 mice were administered with PBS, ALC-0315 LNPs, or crosslinked AP23C5_25+ cLNPs (based on ALC-0315 LNPs) loaded with mOVA via *i.m.* injection (10Lμg mOVA per injection). Mice were sacrificed 7 days after the final injection, and their splenocytes were isolated for analysis **(j)**. The percentages of OVA-specific CD8 T cells (B220^-^ CD3^+^CD8^+^OVA^+^ cells) **(k)**. Splenocytes were restimulated in vitro with OVA and SIINFEKL peptide (100LμgLml^−1^ OVA and 2LμgLml^−1^ SIINFEKL) for 6Lh and assessed via flow cytometry and intracellular cytokine staining to determine the percentages of CD3^+^CD8^+^IFN-γ^+^ **(l)** and CD3^+^CD8^+^TNF-α^+^ **(m)**. **n.** The number of B220^-^CD3^+^CD8^+^ OVA-specific T cells in the spleen was analyzed through flow cytometry. **o.** Frequency of IFN-γ-producing cells in SFU among restimulated splenocytes, assessed via ELISpot. **p**–**r.** Titers of OVA-specific IgG **(p)**, IgG1 **(q)**, and IgG2c **(r)** antibodies in blood serum on Day 21, determined by ELISA. Data are from n = 7 (**a**–**r**) biologically independent samples. For boxplots, the box extends from the 25^th^ to the 75^th^ percentiles, and the line in the middle of the box is plotted at the median. Data were analyzed using one-way ANOVA for **b**–**i**, **k**–**r**. NS: *P* > 0.05, \**P*L<L0.05, \*\**P*L<L0.01, \*\*\**P*L<L0.001, \*\*\*\**P*L<L0.0001.

To extend the applicability of crosslinking strategy, we adapted it for the ALC-0315 LNP formulation, which was used in the BioNTech/Pfzier COVID-19 vaccine. Following the same vaccination regimen, mice were administered i.m. injections of mOVA-loaded ALC-0315 cLNPs on Days 0, 7, and 14, with mOVA-loaded uncrosslinked ALC-0315 LNPs serving as controls (**Fig. 5j**). A marked enhancement in OVA^+^ CD8^+^ T cell frequencies and numbers were observed in the spleens of treated mice, approximately 1.7-fold higher than those in mice receiving the original uncrosslinked ALC-0315 LNPs (**Fig. 5k**, **n****, Supplementary Fig. S20**). Mice treated with the ALC-0315 cLNPs also exhibited increased frequencies of CD3^+^ CD8^+^ IFN-γ^+^ and CD3^+^ CD8^+^ TNF-α^+^cells, approximately 1.9-fold and 1.8-fold respectively, compared to the uncrosslinked control group (**Fig. 5l** and **m**, **Supplementary Figs. S21** and **S22**). Furthermore, ELISpot analysis showed that the crosslinked LNP formulation elicited a ∼1.5-fold increase in the number of IFN-γ-secreting cells compared to ALC-0315 LNPs (**Fig. 5o**, **Supplementary Fig. S23**). In addition, the crosslinked LNPs generated a stronger OVA-specific IgG response, with a ∼258-fold increase in total IgG levels (**Fig. 5p**). These results illustrated the effectiveness of cLNPs in enhancing antigen-specific cellular and humoral immune responses compared to the original ALC-0315 LNP formulation. Given that the crosslinking strategy improved the antigen-specific immune responses for two distinct clinically used LNP formulations, it has the potential to be used across different LNP platforms, enhancing the delivery and efficacy of a variety of mRNA-based vaccines and therapeutics.

Given the enhanced stability of crosslinked mRNA LNP formulations following lyophilization, we next sought to assess the immune activation capability of the lyophilized mOVA-loaded AP23C5_25+ cLNPs, and compare with the lyophilized uncrosslinked mOVA SM-102 LNPs as a control (**Supplementary Fig. S24a**). Mice received *i.m.* vaccinations on Days 0, 7, and 14; and lymphocytes isolated from the spleens on Day 21 were subsequently analyzed to evaluate cellular immune activation. As shown in **Supplementary Fig. S24b**, treatment with lyophilized mRNA cLNPs not only significantly increased the frequency of OVA^+^CD8^+^ T cells in the spleen by approximately 3.9-fold, compared to the lyophilized uncrosslinked SM-102 LNPs, but also achieved frequencies comparable to those observed with freshly prepared SM-102 LNPs. Further analyses revealed that the frequencies of CD3^+^CD8^+^IFN-γ^+^ (**Supplementary Fig. S24c**) and CD3^+^CD8^+^TNF-α^+^ (**Supplementary Fig. S24d**) cells were elevated by approximately 4.8-fold and 3.4-fold, respectively, compared to the control group treated with lyophilized SM-102 LNPs. The humoral immune responses were also enhanced in the lyophilized cLNP group, displaying approximately 15,000-, 322-, and 7,700-fold higher OVA-specific IgG (**Supplementary Fig. S24e**), IgG1 (**Supplementary Fig. S24f**), and IgG2C (**Supplementary Fig. S24g**) titers, respectively. These results demonstrate the potential of mRNA cLNP formulations as a superior platform for efficient mRNA vaccine delivery and robust immune activation, even under conditions where lyophilization of the particles is necessary.

### Enhanced tumor suppression by mRNA cLNPs

To investigate the prophylactic efficacy of the mRNA cLNP formulations as cancer vaccines, we employed B16-OVA melanoma murine tumor model. C57BL/6 mice were immunized with cLNPs encapsulating 10 μg of mOVA mRNA on Days 0, 7, and 14, followed by a subcutaneous (*s.c.*) challenge with 1 × 10^6^ B16-OVA cells in the right flank on Day 28 (**Fig. 6a**). cLNP formulations demonstrated superior tumor growth inhibition and increased survival compared to the uncrosslinked formulation (**Fig. 6b**–**d**). Mice immunized with crosslinked versions of SM-102 and ALC-0315 LNPs showed median survival times extended to 23.5 and 23 days, respectively, compared to 19 and 19.5 days with the original formulations. Tumor occurrence was delayed until Day 18 in mice treated with cLNPs, whereas tumors were evident by Day 10 in those receiving the uncrosslinked LNPs.

**Figure 6.**
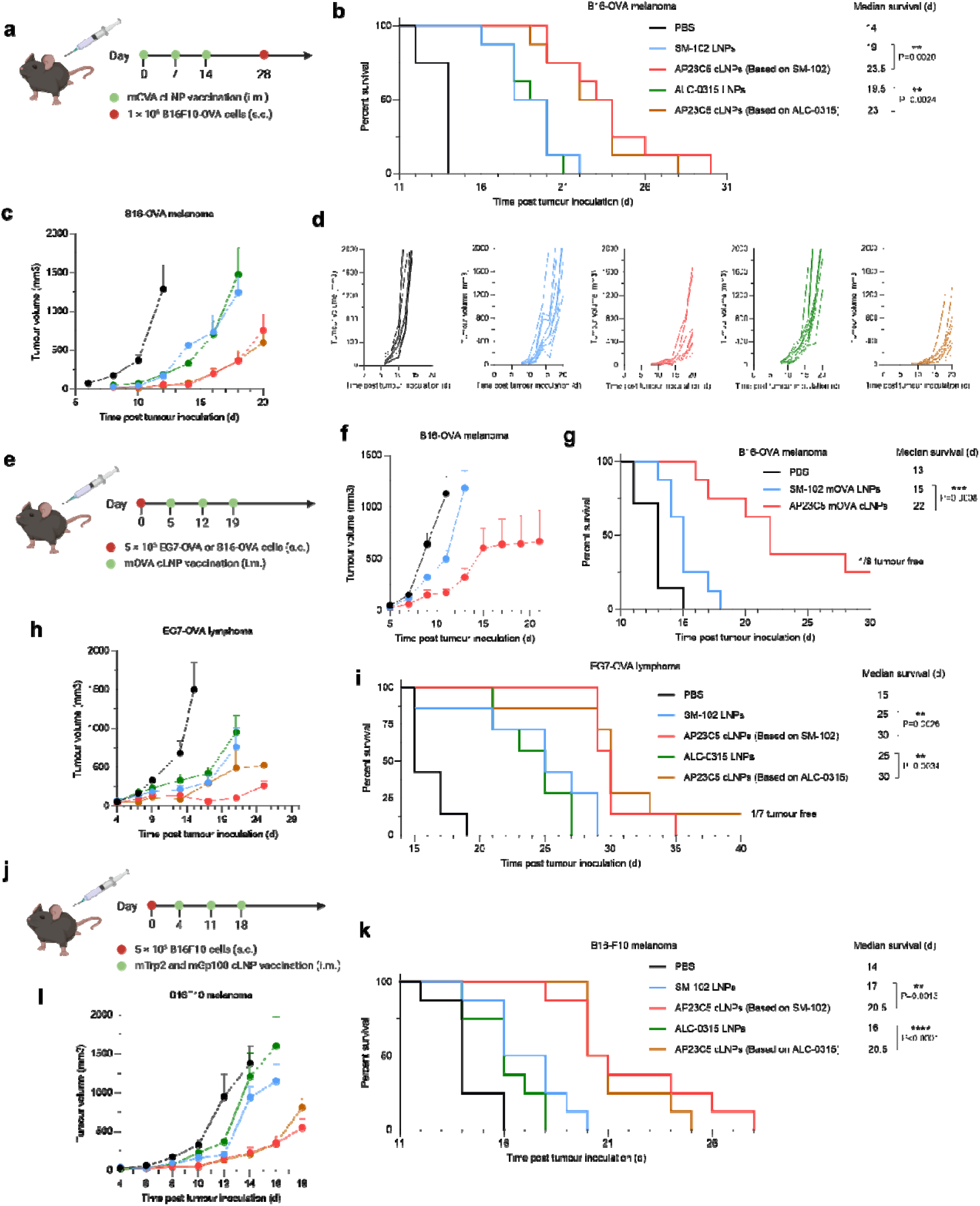
Antitumor efficacy of mRNA cLNP formulations as prophylactic or therapeutic vaccines. a–d. Schematic and results of a prophylactic vaccination model using OVA-expressing melanoma in C57BL/6 mice. Mice received three *i.m.* injections (10 μg mOVA per injection), one week apart, of PBS, mOVA-loaded SM-102 LNPs, AP23C5_25+ SM-102 cLNPs, ALC-0315 LNPs, or AP23C5_25+ ALC-0315 cLNPs. Following these injections, mice were *s.c.* inoculated with OVA-expressing melanoma (B16-OVA) cells **(a)**. The results presented include survival curves **(b)**, average tumor volume **(c)**, and individual tumor volumes **(d)** over time. **e.** Schematic and results of a therapeutic vaccination model for B16-OVA and EG7-OVA in C57BL/6 mice. Mice were *s.c.* inoculated with B16-OVA cells and subsequently received three *i.m*. injections, one week apart, of PBS, mOVA-loaded SM-102 LNPs, or AP23C5_25+ SM-102 cLNPs. The results displayed include average tumor volumes **(f)** and survival curves **(g)**. Mice were *s.c.* inoculated with EG7-OVA cells and subsequently received three *i.m.* injections, one week apart, of PBS, mOVA-loaded SM-102 LNPs, AP23C5_25+ SM-102 cLNPs, ALC-0315 LNPs, or AP23C5_25+ ALC-0315 cLNPs. The results displayed include average tumor volumes **(h)** and survival curves **(i)**. **j–l**, Schematic and results of a therapeutic vaccine against melanoma-associated antigens for melanoma in C57BL/6 mice. Mice were inoculated *s.c.* with B16F10 cells and then given three *i.m.* injections, one week apart, of PBS or SM-102 LNPs, AP23C5_25+ SM-102 cLNPs, ALC-0315 LNPs, or AP23C5_25+ ALC-0315 cLNPs loaded with mRNA encoding Trp2 (mTrp) or Gp100 (mGp100) (10 μg total mRNA per injection). The results displayed include survival curves **(k)** and average tumor volumes **(l)**. Data represent meanL±Ls.e.m. with nL=L8 (**b-d, f-g, k-l**), n = 7 (**h-i**) biologically independent samples. Survival curves were compared using the log-rank Mantel-Cox test. NS: *P* > 0.05, \**P* L<L0.05, \*\**P* L<L0.01, \*\*\**P* L<L0.001, \*\*\*\**P* L<L0.0001.

Next, we evaluated the therapeutic efficacy of the crosslinked LNPs in B16 and EG7 tumor models, using OVA as well as melanoma-associated antigens Trp2 and Gp100. Mice received tumor cells on Day 0 and were treated with cLNPs containing 10 μg of mOVA mRNA on Days 5, 12, and 19 (**Fig. 6e**). The cLNPs led to a marked tumor suppression in the B16-OVA model, improving the median survival from 15 to 22 days, with one mouse remaining tumor-free at Day 30 (**Fig. 6g**, **h****, Supplementary Fig. S25e**). Similar results were observed in the EG7-OVA model (**Fig. 6i**, **j**), where cLNPs significantly prolonged survival to 30 days, outperforming the uncrosslinked controls. Remarkably, among the mice treated with the ALC-0315 cLNPs, one in seven displayed no detectable tumors 30 days post-tumor inoculation.

Subsequent experiments employed clinically relevant tumor antigens Trp2 and Gp100 in the same tumor model. On Day 0, 5 × 10^5^ B16F10 cells were subcutaneously injected into the right flank of C57BL/6 mice. Mice were vaccinated on Days 4, 11, and 18 with cLNPs carrying 10 μg of mRNA encoding Trp2 and Gp100 (**Fig. 6k**). The cLNPs loaded with these antigens also demonstrated significant anti-tumor activity (*P* < 0.01), extending median survival times to 20 days compared to the uncrosslinked SM-102 and ALC-0315 LNPs (**Fig. 6l**, **m****, Supplementary Fig. S26**). These results highlight that cLNP formulations enhance the delivery efficiency of functional mRNAs, inducing more effective anti-tumor responses and improving cancer therapies.

## Discussion

Ensuring stability has been a challenge for LNP-based mRNA delivery. Commonly explored stabilization techniques, including optimization of lipid components like ionizable lipids^35, 36^, cholesterol derivatives^37^, and phospholipids^38^, buffer modifications^15^, and inclusion of cryoprotectants like trehalose^39^, have not generated robust and generalizable outcomes. On the other hand, crosslinking via covalent or physical bonds has been intensively studied to enhance the stability, release mechanisms, and targeting efficiency of polymer-based nanotherapeutics. For example, photo-crosslinking has been employed to enhance the stability of bioreducible polymer nanoparticles^40^, thereby improving their functionality as a systemic delivery carrier for siRNA in cancer therapy. Similarly, a series of core crosslinked star polymers with various compositions have been shown to increase the delivery efficiency of siRNA to pancreatic cancer cells^41^. Despite extensive research on crosslink-based stabilization methods for polymer particles, their application in mRNA lipid nanoparticles remains largely unexplored.

For LNP crosslinking, the selection of crosslinking schemes – either covalent or physical crosslinking–requires careful titration of the crosslinking structure and degree, which can influence not only the stability, but also the function and efficacy of the cLNPs. Compared with physical crosslinking methods, covalent crosslinking offers significant advantages due to the higher stability to maintain the structural integrity of LNPs, affording a higher degree of resistance to environmental changes such as temperature fluctuations, which is crucial for long-term storage and transport. Moreover, the design of biological environment-sensitive covalent crosslinks can aid in achieving a triggered or controlled release of genetic payload once the LNPs reach the cytosol, for therapeutic applications that benefit from enhanced and prolonged transgene expression.

In this study, we developed an effective, easy-to-adapt, and generalizable crosslinking approach using cholesterol-derived agents that form acid-sensitive cleavable bonds, which substantially enhanced the structural integrity of LNPs. This enhanced stability was evident from the reduced release of payloads from cLNPs under the dextran sulfate challenge. The choice of cholesterol in our crosslinking strategy is based on it as a ubiquitous component in all LNPs and informed by its essential role in stabilizing lipid bilayers, extending circulation times, and enhancing cellular uptake—key factors for the successful delivery of LNPs^42, 43^. Leveraging the relatively high molar ratio of cholesterol in LNP formulations, our strategy strengthens LNP stability and functionality without compromising biocompatibility and efficiency of the delivery.

Optimizing the crosslinking structure within cLNPs requires consideration of achieving the correct balance of crosslinking degree—a critical factor influencing both structural integrity and the efficacy of mRNA delivery. A higher crosslinking degree generally enhances the structural integrity of LNPs, making them more stable under production, storage, and physiological conditions. This stability will confer protection to LNPs from premature degradation during circulation, ensuring that they reach their target tissues or cells intact. However, in cases where the crosslinking degree is so high that the LNPs are too stable, the release of mRNA could be negatively impacted once the LNPs enter their target cells, slowing down the necessary disintegration for effective mRNA release into the cytosol. An optimal crosslinking degree ensures that LNPs are sufficiently stable to protect their payload during transit while also effectively releases the mRNA cargo in cytosol. The optimal crosslinking structure and degree require experimental fine-tuning through exploration within a large library of crosslinking conditions as we have shown in this study. This optimization ensures that the mRNA cLNPs can fulfill their role as effective delivery vehicles for mRNA, maximizing therapeutic outcomes while minimizing potential side effects.

By optimizing the crosslinking degree and structure of the cLNPs through *in vitr*o and *in vivo* screening, we observed a marked improvement in mRNA delivery, achieving higher transfection efficiencies compared to original uncrosslinked formulations. Importantly, the design of the cleavable crosslinking structure played a crucial role in facilitating endosomal escape, a critical step for effective intracellular mRNA delivery. Additionally, the crosslinked structures maintained their physical properties even after lyophilization and freeze-thaw cycles, underscoring their robustness and suitability for handling and storage. We demonstrated the effectiveness of this crosslinking approach by applying it to both Moderna and Pfizer commercial LNP formulations and a previously reported custom-tailored LNP formulation^21^, showing enhanced immune responses against cLNPs loaded with antigen-expressing mRNA, as evident by the increased T-cell activation and cytokine production, and more importantly anti-tumor efficacy. These findings highlight the dual benefits of our crosslinking strategy, which not only improve the physical stability of the LNPs but also enhances their functionality as an mRNA delivery system. The improved stability and intracellular delivery of mRNA facilitated by our crosslinking strategy make these cLNPs promising candidates for a variety of mRNA-based therapies and vaccines, advancing the potential of LNP-mediated gene delivery for clinical applications.

In summary, we have developed a post-assembly crosslinking strategy for stabilizing mRNA LNPs by incorporating a series of cholesterol derivatives and crosslinkers. The mRNA cLNP formulations demonstrated improved physical stability under physiological and lyophilization conditions, enhanced cellular uptake and endosomal escape, and elevated transfection efficiency compared to uncrosslinked formulations. In addition, these cLNPs extended *in vivo* mRNA expression duration and induced robust antigen-specific immune responses. The versatility of the strategy was further showcased by its successful adaptation in different LNP formulations, establishing crosslinked mRNA LNPs as a promising platform for mRNA-based therapeutics and vaccines, with the potential for sustained gene delivery.

### Methods Materials

Cholesteryl hemisuccinate (CHEMS), 1,1,1-tris(hydroxymethyl)ethane, 2,2-dimethoxypropane, 5-hydroxyisophthalaldehyde, 4-hydroxy-benzaldehyde, p-toluenesulfonic acid monohydrate (p-TsOH), 5,5-dimethyl-1,3-cyclohexanedione, 1-(3-dimethyl aminopropyl)-3-ethylcarbodiimide hydrochloride (EDC), K_2_CO_3_, triethylamine (TEA), 4-dimethyl aminopyridine (DMAP), Dess-Martin periodinane (DMP), Lys-Lys-Lys, 1,4-diamino-butane, butane-1,4-diol, dichloromethane, methanol, n-hexane, ethyl acetate (EtOAc), and cholesterol were purchased from Sigma-Aldrich. Cyclohexane-1,4-diyldimethanamine, 1,8-diamino-octane, 1,12-diamino-dodecane were purchased from Ambeed. PEG1 diamine, PEG2 diamine, PEG4 diamine, and PEG8 diamine were purchased from BroadPharm Biotechnology. PEG23 diamine and PEG45 diamine were purchased from Biopharma PEG Biotechnology. SM-102, ALC-0315, and DLin-MC3-DMA were purchased from MedKoo Biosciences. DSPC, DOPE, and DMG-PEG2000 were obtained from Avanti Polar Lipids. B16F10 cells (CRL-6475) were purchased from the American Type Culture Collection (ATCC). DC2.4 cells and B16-OVA (expressing model antigen, OVA, with a transmembrane domain) were kindly provided by the lab of Prof. Jonathan Schneck. Reporter lysis buffer and luciferin assay solution were purchased from Promega. All mRNA was purchased from Hongene Biotechnology. D-luciferin was purchased from Gold Biotechnology.

### Cholesterol-derived crosslinker (CDCL) synthesis

Details on the synthesis and characterization of the CDCL are provided in the Supplementary Information. ^1^H NMR spectra were recorded using a Bruker 400-MHz NMR spectrometer.

### Biological reagents

Antibodies used in this study are: APC-Cy7, Brilliant Violet 750 anti-mouse CD8a (BioLegend, 100714,747134); Brilliant Violet 421 anti-OVA SIIGFEKL (Tetramer) (NIH tetramer core facility, N/A); APC, PE, Brilliant Violet 421 anti-mouse CD3 (BioLegend, 100236,100206, 100228); Live/Dead Fix Aqua (Thermo Fisher, L34957); FITC anti-mouse CD44 (BioLegend, 103006); PE anti-mouse CD62L (BioLegend, 161204); FITC, Brilliant Violet 421 anti-mouse anti-mouse CD45 (BioLegend, 103108,103134); APC-Cy7 anti-mouse CD31 (BioLegend, 102440); Brilliant Violet 605,APC anti-mouse CD326 (BioLegend, 118227, 118214) ;FITC anti-mouse CD8 (BioLegend, 100706); Live/Dead Fix Near IR (780) (Thermo Fisher, L10119); PE-Cy7 anti-mouse CD45R (BioLegend, 103222); Alexa Fluor 700 anti-mouse CD69 (BioLegend, 104539); Alexa Fluor 700 anti-mouse TNF-α (BioLegend, 506338); PE-Cy7 anti-mouse CD4 (BioLegend, 100422); Brilliant Violet 421 anti-mouse CD86 (BioLegend, 105032); Brilliant Violet 750 anti-mouse CD11c (BioLegend, 117357).

### Cell culture and high-throughput screening for transfection studies

For monolayer culture studies, DC2.4 cells were seeded into 96-well plates at a cell density of 10,000 cells per well 1 day before transfection. LNPs were pipetted into RPMI medium at a final concentration of 1 μg mL^−1^ of mRNA. For example, 5 μL of cLNPs suspension at 20 μg mL^−1^ of mRNA was pipetted into the 100 μL culture medium in each well. The transgene expression was analyzed following 24 h incubation. When characterizing luciferase as the reporter, cells were lysed by reporter lysis buffer (Promega) using two freeze-thaw cycles, with the lysate characterized by a luminometer upon the addition of luciferin assay solution (Promega) against a standard curve generated using luciferase samples (Promega).

### Synthesis and characterization of mRNA cLNPs

The preparation process of AP23C5_25+ cLNPs is described as an example. The cLNPs were synthesized by adding an organic phase containing the lipids to an aqueous phase containing the mRNAs in a 96-well plate and further adjusting the pH to 7.4 for high-throughput screening. To prepare the organic phase, a mixture of SM-102, cholesterol, CDCL5, DSPC and DMG-PEG2000 were dissolved in ethanol at a molar ratio of 50:38.5:9.63:10:1.5. To prepare the aqueous phase, corresponding mRNA (fLuc mRNA, mCherry mRNA, OVA mRNA, Trp2 mRNA or Gp100 mRNA) was prepared in 25 mM magnesium acetate buffer (pH 4.0, Fisher) containing PEG23 diamine. All mRNA samples were stored at −80 °C and thawed on ice before

use. For *in vitro* screening, cLNPs were incubated with cells without dialysis. For larger-scale cLNPs production, the aqueous and ethanol phases prepared were mixed at a 3:1 ratio in a T-junction device using syringe pumps, further adjusted to pH 7.4, and then purified by dialysis against PBS using a 3.5-kDa molecular weight cut-off (MWCO) Pur-A-Lyzer (Sigma-Aldrich) at 4 °C for 24 h and stored at 4 °C before injection. The size, polydispersity index and zeta potentials of cLNPs were measured using dynamic light scattering (ZetaPALS, Brookhaven Instruments). Diameters are reported as the intensity mean average. The morphology of the cLNPs was characterized by a cryo-electron microscope (Glacios, Thermo Fisher).

A Quant-iT RiboGreen assay (ThermoFisher, R11490) was utilized to quantify mRNA concentrations and assess the encapsulation efficiency (EE) of crosslinked mRNA LNPs. Standard curves were prepared in duplicate on a black 96-well plate with mRNA concentrations ranging from 0.1 to 1.0 µg/mL in each well. Samples of LNPs were added in triplicate at a consistent LNP to DI water ratio of 1:20. Following this, 60 µL of RiboGreen solution—a 200-fold dilution of RiboGreen dye in water—was added to each well. Fluorescence measurements were taken using a BioTek Synergy H1 plate reader (excitation: 480 nm, emission: 520 nm) to determine mRNA levels in intact LNPs. Following the initial reading, 10% w/v Triton-X was added to each well with shaking to disrupt the cLNPs structure and release the encapsulated mRNA. Similar to the first reading, 60 µL of RiboGreen solution was added to the wells. Fluorescence readings were measured to determine the total mRNA content post-release. The measured fluorescence signals before and after mRNA release were converted to mRNA concentration using the standard curves. The EE% of the cLNPs were calculated using the following equation.

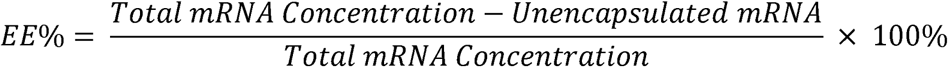

### Lyophilization and stability analysis

Transfer the crosslinked mRNA LNPs solution into a 96-well plate, then freeze at −45°C and lyophilize with a freeze-dryer (Triad). Reconstitute the lyophilized mRNA cLNPs in PBS and assess their size and polydispersity index using dynamic light scattering (ZetaPALS, Brookhaven Instruments); diameters are reported as intensity-weighted means. Following the same protocol, evaluate transfection efficiency. Briefly, seed DC2.4 cells in a 96-well plate at 10,000 cells per well one day prior to transfection. Introduce the lyophilized cLNPs into RPMI medium, achieving a final concentration of 1 μg/mL mRNA. After 24 h of incubation, analyze the expression of the transgene. When characterizing luciferase as the reporter, cells were lysed by reporter lysis buffer (Promega) using two freeze–thaw cycles, with the lysate characterized by a luminometer upon addition of luciferin assay solution (Promega) against a standard curve generated using luciferase samples (Promega).

### Assessment of cellular uptake and endosomal escape efficiency in mRNA cLNPs

The labeled mRNA cLNPs were prepared using Cy5-labeled mRNA (TriLink). DC2.4 cells were seeded at a density of 10,000 per well in 96-well plates, and transfected with Cy5-labeled cLNPs at a dose equivalent to 0.1 µg per well of mRNA in media. Cells were washed and stained with LIVE/DEAD Fixable Aqua Dead Cell Stain Kit (Thermo Fisher Scientific) at 4 h after transfection, and analyzed by flow cytometry as described above; the data were processed using FlowJo 10.

B16-F10 cell line expressing GFP-coupled galectin-8 (GFP-Gal8) was obtained by transfection using plasmids encoding Super PiggyBac Transposase (System Biosciences) and Piggybac transposon-GFP-Gal8 (Addgene plasmid #127191) and a poly(β-amino ester) carrier, and then sorted by an SH800 cell sorter (Sony) twice. B16-GFP-Gal8 cells were seeded at a density of 5,000 per well in 96-well plates and transfected with crosslinked mRNA LNPs at a dose equivalent to 0.1 µg per well of mRNA in media. At 4 h after transfection, cells were washed with PBS three times, fixed with 4% paraformaldehyde and stained with Hoechst 33342. The plates were then analyzed by a CellInsight CX7 HCA platform (Thermo Fisher Scientific). A total of 10 fields were analyzed inside each well of the plates, and the well-averaged result was generated by averaging all the cells in all the fields. For confocal laser scanning microscopy imaging, transfected cells were fixed with 10% paraformaldehyde for 10 min at room temperature, washed with PBS and Hoechst nuclear stained (1:1,000 in PBS) before imaging on an Apotome microscope (Zeiss).

### Animals and primary cells

All animal procedures were conducted in accordance with protocols approved by the Johns Hopkins Institutional Animal Care and Use Committee (Protocol #MO23E31). C57BL/6 mice (male and female), aged 6–8 weeks, were obtained from the Jackson Laboratory. The mice had free access to pelleted feed and water, with the feed typically containing 5% fiber, 20% protein, and 5–10% fat. On average, the mice consumed 4–5 g of pelleted feed (120 g per kg body weight) and drank 3–5 mL of water (150 mL per kg body weight) daily. The temperature in the mouse rooms was maintained between 18–26 °C (64–79 °F) with 30–70% relative humidity, ensuring at least 10 air changes per hour. The mice were housed in standard shoebox cages with corncob bedding.

The LNPs were given through *i.m.* (right quadriceps) injection at a predetermined dose per mouse. The LNP suspensions were concentrated to 200 μg/mL for *i.m.* injection of mRNA by an Amicon Ultra-2 centrifugal filter unit with an MWCO of 100 kDa. The D-luciferin solution was given through intraperitoneal (*i.p.*) injection (lower quadrant of abdomen) at a predetermined dose per mouse.

### Tissue processing and cell isolation

For isolation of lymphocytes from the spleen, harvested spleens were placed onto 40-μm cell strainers. The spleens were mechanically digested through the cell strainers with the back of a 3-mL syringe plunger in a lymphocyte separation medium (PromoCell). The resulting single-cell cell suspensions were transferred to sterile 15-mL tubes, and 1 mL of RPMI-1640 media was slowly added to each tube to form visible layers. The tubes were then centrifuged at 800 ×g for 25 min at 4 °C without brake. Following centrifugation, the middle layer containing lymphocytes was carefully collected and transferred to fresh 15-mL tubes, and 5 mL of RPMI-1640 media was added and mixed thoroughly. The cells were pelleted by centrifugation at 300 ×g for 5 min at 4 °C. The pellet was then resuspended in 3 mL of RPMI-1640 media and centrifuged again at 300 ×g for 5 min at 4 °C. Isolated lymphocytes were suspended in RPMI-1640 media for subsequent analyses.

### Antibodies, cell isolation and staining for flow cytometry

All antibodies were diluted at a ratio of 1:100 before use. LIVE/DEAD fixable dead cell stain kits were used to determine the viability of cells. CellTrace Violet cell proliferation kit (ThermoFisher, C34571) was used to stain and monitor OT-I cells in vivo. eBioscience Foxp3/Transcription Factor Staining buffer set (ThermoFisher, 00-5523-00) was used for intracellular staining.

Isolated cells from the tissues, as described in the previous section, were resuspended and pelleted in 100 µL of antibodies diluted in flow cytometry staining buffer obtained from eBioscience™. The cells were then incubated on ice in the dark for 1 h. After the incubation period, the stained cells were washed twice with PBS and subsequently resuspended in 200 µL of eBioscience™ flow cytometry staining buffer for flow cytometry analysis. Flow data was acquired using an Attune NXT flow cytometer and analyzed with FlowJo software v.10.

### Enzyme-linked immunosorbent spot (ELISpot) assay

For the ELISpot assay, multiscreen filter plates (Millipore-Sigma, S2EM004M99) were coated with antibodies targeting IFN-γ (BD Biosciences, 551881) and subsequently blocked according to the manufacturer’s instructions. A total of 1×106 isolated lymphocytes were then added to each well and stimulated with SIINFEKL peptide at a concentration of 2 μg/mL for 18 h. The spots were detected using a mouse IFN-γ detection antibody (BD Biosciences, 551881), followed by an incubation with streptavidin-HRP (BD Biosciences, 557630) and AEC substrate (BD Biosciences, 551951). The plates were then forwarded to the SKCCC Immune Monitoring Core for further analysis.

### Enzyme-linked immunosorbent assay (ELISA)

For serum antibody detection, 100 μL of blood sample was drawn from the tail vein of immunized mice on the day of sacrifice. Levels of antigen-specific IgG, IgG1, and IgG2c in the serum were measured by ELISA. Flat-bottomed 96-well plates (Nunc) were precoated with OVA protein at a concentration of 2 μg protein per well in 100 mM carbonate buffer (pH 9.6) at 4°C overnight, which were then blocked at room temperature for 2 h with blocking buffer, which consisted of 10% BSA in PBS. The serum obtained from mice was first diluted 30 times in the blocking buffer, followed by threefold serial dilutions. After blocking was done, the plates were washed with PBS-T (PBS containing 0.05% Tween), and the diluted serum samples were then added to the wells and incubated at 4 °C overnight. Horseradish peroxidase-conjugated goat anti-mouse IgG, IgG1, and IgG2c (Southern Biotech Associates) were used at a dilution of 1:2,000, 1:4,000, and 1:4,000, respectively, in the blocking buffer for labeling. After 1-h antibody incubation at room temperature, the plates were washed with PBS-T, followed by incubation with TMB ELISA substrate solution at room temperature. After a 30-min incubation, 50 µL of 4 N sulfuric acid was added to the wells to quench the reaction. The plates were then read at a wavelength of 450 nm with a plate reader. A sample was considered positive if its absorbance was twice as much as or higher than the absorbance of the negative control.

### Immunization and tumor therapy experiments

Mice aged 6–8 weeks were injected subcutaneously with B16-OVA cells (1 × 10^6^ in prophylactic and 5 × 10^5^ in therapeutic studies) or 5 × 10^5^ B16F10 melanoma cells or 5 × 10^5^ EG7-OVA lymphoma cells into the right flank. In therapeutic studies, vaccinations began when tumor sizes were less than 50 mm^3^. Animals were immunized by *i.m.* injection of different LNP formulations containing 10 μg OVA mRNA, mTrp2, or mGp100 as described in the main text. A total of three doses were given. Tumor growth was measured three times a week using a digital caliper and calculated as 0.5 × length × width × width. Mice were euthanized when the tumor volumes reached 2,000 mm^3^.

### Statistics

Unpaired t-tests were performed when comparing two groups. One-way analysis of variance (ANOVA) and Tukey’s multiple comparisons were performed when comparing more than two groups. Survival curves were compared using log-rank Mantel–Cox test. Statistical analysis was performed using Microsoft Excel and Prism 10 (GraphPad). Statistical analysis was performed using Microsoft Excel and Prism 10 (GraphPad). A difference was considered significant if *P* < 0.05 (\**P* < 0.05, \*\**P* < 0.01, \*\*\**P* < 0.001, \*\*\*\**P* < 0.0001).

## Funding

This study is partially supported by P41EB028239 (H.Q.M.).

## Author contributions

X.L., Y.Z. and H.-Q.M. conceived of and designed this study. H.-Q.M. secured the funding for this study. X.L., Y.Z., C.W., J.L., D.Y., J.K., F.S., J.M., T.X., Y.S., K.D.G., L.C. W.H.T. C.J.E., and S.L, performed the experiments. X.L., Y.Z., C.W., J.L., D.Y., J.K., F.S., T-H.W. and H.-Q.M. participated in data analysis and interpretation. The manuscript was written by X.L., Y.Z. and H.-Q.M., with revisions by C.W. and inputs from all the other authors.

## Competing interests

H.-Q.M., Y.Z., and X.L. are co-inventors of a pending patent application covering the crosslinked LNP formulations described in this paper, filed through and managed by Johns Hopkins Technology Ventures. The other authors declare no competing interests.

## Data and materials availability

All data needed to evaluate the conclusions in the paper are present in the paper and/or the Supplementary Materials.

## Code availability statement

There is no code used in the paper.

## References

1. Huang, X. et al. Nanotechnology-based strategies against SARS-CoV-2 variants. Nat Nanotechnol 17, 1027–1037 (2022).

2. Liu, S. et al. Membrane-destabilizing ionizable phospholipids for organ-selective mRNA delivery and CRISPR-Cas gene editing. Nat Mater 20, 701–710 (2021).

3. Hou, X., Zaks, T., Langer, R. & Dong, Y. Lipid nanoparticles for mRNA delivery. Nat Rev Mater 6, 1078–1094 (2021).

4. Barbier, A.J., Jiang, A.Y., Zhang, P., Wooster, R. & Anderson, D.G. The clinical progress of mRNA vaccines and immunotherapies. Nat Biotechnol 40, 840–854 (2022).

5. Warne, N. et al. Delivering 3 billion doses of Comirnaty in 2021. Nat Biotechnol 41, 183–188 (2023).

6. Baden, L.R. et al. Efficacy and safety of the mRNA-1273 SARS-CoV-2 vaccine. N Engl J Med 384, 403–416 (2021).

7. Polack, F.P. et al. Safety and efficacy of the BNT162b2 mRNA Covid-19 vaccine. N Engl J Med 383, 2603–2615 (2020).

8. Chaudhary, N., Weissman, D. & Whitehead, K.A. mRNA vaccines for infectious diseases: principles, delivery and clinical translation. Nat Rev Drug Discov 20, 817–838 (2021).

9. Liu, C., et al. mRNA-based cancer therapeutics. Nat Rev Cancer 23, 526–543 (2023).

10. Sharma, P., Hoorn, D., Aitha, A., Breier, D. & Peer, D. The immunostimulatory nature of mRNA lipid nanoparticles. Adv Drug Deliv Rev 205, 115175 (2024).

11. Samaridou, E., Heyes, J. & Lutwyche, P. Lipid nanoparticles for nucleic acid delivery: Current perspectives. Adv Drug Deliv Rev 154-155, 37–63 (2020).

12. Chen, S., et al. Nanotechnology-based mRNA vaccines. Nature Reviews Methods Primers 3, 63 (2023).

13. Slezak, A. et al. Therapeutic synthetic and natural materials for immunoengineering. Chem Soc Rev 53, 1789–1822 (2024).

14. Bitounis, D., Jacquinet, E., Rogers, M.A. & Amiji, M.M. Strategies to reduce the risks of mRNA drug and vaccine toxicity. Nat Rev Drug Discov 23, 281–300 (2024).

15. Jiang, A.Y. et al. Combinatorial development of nebulized mRNA delivery formulations for the lungs. Nat Nanotechnol 19, 364–375 (2024).

16. Gupta, A., Andresen, J.L., Manan, R.S. & Langer, R. Nucleic acid delivery for therapeutic applications. Adv Drug Deliv Rev 178, 113834 (2021).

17. Liu, S. et al. Charge-assisted stabilization of lipid nanoparticles enables inhaled mRNA delivery for mucosal vaccination. Nat Commun 15, 9471 (2024).

18. Xue, L. et al. Combinatorial design of siloxane-incorporated lipid nanoparticles augments intracellular processing for tissue-specific mRNA therapeutic delivery. Nat Nanotechnol 20, 132–143 (2025).

19. Zhao, S. et al. Acid-degradable lipid nanoparticles enhance the delivery of mRNA. Nat Nanotechnol 19, 1702–1711 (2024).

20. Li, B. et al. Combinatorial design of nanoparticles for pulmonary mRNA delivery and genome editing. Nat Biotechnol 41, 1410–1415 (2023).

21. Zhu, Y. et al. Screening for lipid nanoparticles that modulate the immune activity of helper T cells towards enhanced antitumour activity. Nat Biomed Eng 8, 544–560 (2024).

22. Rhym, L.H., Manan, R.S., Koller, A., Stephanie, G. & Anderson, D.G. Peptide-encoding mRNA barcodes for the high-throughput in vivo screening of libraries of lipid nanoparticles for mRNA delivery. Nat Biomed Eng 7, 901–910 (2023).

23. Lokugamage, M.P. et al. Optimization of lipid nanoparticles for the delivery of nebulized therapeutic mRNA to the lungs. Nat Biomed Eng 5, 1059–1068 (2021).

24. Lian, X. et al. Bone-marrow-homing lipid nanoparticles for genome editing in diseased and malignant haematopoietic stem cells. Nat Nanotechnol 19, 1409–1417 (2024).

25. Brown, D.W. et al. Safe and effective in vivo delivery of DNA and RNA using proteolipid vehicles. Cell 187, 5357–5375.e24 (2024).

26. Cheng, Q. et al. Selective organ targeting (SORT) nanoparticles for tissue-specific mRNA delivery and CRISPR-Cas gene editing. Nat Nanotechnol 15, 313–320 (2020).

27. Mitchell, M.J. et al. Engineering precision nanoparticles for drug delivery. Nat Rev Drug Discov 20, 101–124 (2021).

28. Li, Y., Xiao, K., Zhu, W., Deng, W. & Lam, K.S. Stimuli-responsive cross-linked micelles for on-demand drug delivery against cancers. Adv Drug Deliv Rev 66, 58–73 (2014).

29. Shae, D. et al. Endosomolytic polymersomes increase the activity of cyclic dinucleotide STING agonists to enhance cancer immunotherapy. Nat Nanotechnol 14, 269–278 (2019).

30. Christensen, P.R., Scheuermann, A.M., Loeffler, K.E. & Helms, B.A. Closed-loop recycling of plastics enabled by dynamic covalent diketoenamine bonds. Nat Chem 11, 442–448 (2019).

31. Li, S. et al. Payload distribution and capacity of mRNA lipid nanoparticles. Nat Commun 13, 5561 (2022).

32. Chatterjee, S., Kon, E., Sharma, P. & Peer, D. Endosomal escape: A bottleneck for LNP-mediated therapeutics. Proc Natl Acad Sci U S A 121, e2307800120 (2024).

33. Zheng, L., Bandara, S.R., Tan, Z. & Leal, C. Lipid nanoparticle topology regulates endosomal escape and delivery of RNA to the cytoplasm. Proc Natl Acad Sci U S A 120, e2301067120 (2023).

34. Rui, Y. et al. High-throughput and high-content bioassay enables tuning of polyester nanoparticles for cellular uptake, endosomal escape, and systemic in vivo delivery of mRNA. Sci Adv 8, eabk2855 (2022).

35. Packer, M., Gyawali, D., Yerabolu, R., Schariter, J. & White, P. A novel mechanism for the loss of mRNA activity in lipid nanoparticle delivery systems. Nat Commun 12, 6777 (2021).

36. Hashiba, K. et al. Overcoming thermostability challenges in mRNA-lipid nanoparticle systems with piperidine-based ionizable lipids. Commun Biol 7, 556 (2024).

37. Zhang, X., Barraza, K.M. & Beauchamp, J.L. Cholesterol provides nonsacrificial protection of membrane lipids from chemical damage at air-water interface. Proc Natl Acad Sci U S A 115, 3255–3260 (2018).

38. Kafle, U., Truong, H.Q., Nguyen, C.T.G. & Meng, F. Development of thermally stable mRNA-LNP delivery systems: Current progress and future prospects. Mol Pharm 21, 5944–5959 (2024).

39. Kafetzis, K.N. et al. The effect of cryoprotectants and storage conditions on the transfection efficiency, stability, and safety of lipid-based nanoparticles for mRNA and DNA delivery. Adv Healthc Mater 12, e2203022 (2023).

40. Karlsson, J. et al. Photocrosslinked bioreducible polymeric nanoparticles for enhanced systemic siRNA delivery as cancer therapy. Adv Funct Mater 31, 2009768 (2021).

41. Teo, J. et al. A rationally optimized nanoparticle system for the delivery of RNA interference therapeutics into pancreatic tumors in vivo. Biomacromolecules 17, 2337–2351 (2016).

42. Back, P.I. et al. Immune implications of cholesterol-containing lipid nanoparticles. ACS Nano 18, 28480–28501 (2024).

43. Patel, S. et al. Naturally-occurring cholesterol analogues in lipid nanoparticles induce polymorphic shape and enhance intracellular delivery of mRNA. Nat Commun 11, 983 (2020).

