## Supporting Information for "Crosslinking of Lipid Nanoparticles Enhances the Delivery Efficiency and Efficacy of mRNA Vaccines"

### Supplementary Figures

**Scheme 1** Synthesis of compound CDCL1.

**Scheme 2** Synthesis of compound CDCL2.

**Scheme 3** Synthesis of compound CDCL3.

**Scheme 4** Synthesis of compound CDCL4.

**Scheme 5** Synthesis of compound CDCL5.

**Scheme 6** Synthesis of compound CHNH.

**Figure 1** Representative $^{1}$H-NMR spectrum of CDCL1.

**Figure 2** Representative $^{1}$H-NMR spectrum of CDCL2.

**Figure 3** Representative $^{1}$H-NMR spectrum of CDCL3.

**Figure 4** Representative $^{1}$H-NMR spectrum of CDCL4.

**Figure 5** Representative $^{1}$H-NMR spectrum of CDCL5.

**Figure 6** Representative $^{13}$C NMR spectrum of CDCL5.

**Figure 7** *In vitro* evaluation of top-performing crosslinking candidates in SM-102 LNPs for transfection and maturation of DCs.

**Figure 8** *In vitro* evaluation of top-performing crosslinking candidates in ALC-0315 LNPs or C10 LNPs for transfection and maturation of DCs.

**Figure 9** The geometric means of TMR signal (indicator of relative helper lipid content) intensity of mRNA loaded LNPs.

**Figure 10** The mRNA payload distribution profiles of SM-102 LNP or A2C5_25+ cLNP formulations before and after dialysis.

**Figure 11** Representative $^{1}$H-NMR spectrum of CHNH.

**Figure 12** H&E staining of major organs of mice treated with SM-102 LNPs or AP23C5_25+ LNPs for in vivo biosafety.

**Figure 13** *In vivo* transfection efficiency of crosslinked vs. uncrosslinked ALC-0315 LNPs.

**Figure 14** *In vivo* transfection efficiency of crosslinked vs. uncrosslinked C10 LNPs.

**Figure 15** Gating strategy for flow cytometry plots for antigen (OVA)-specific T cell responses in the spleen after vaccination.

**Figure 16** Representative flow cytometry plots for determining OVA-specific CD8$^{+}$ T cells in the spleen are shown by the SM-102 mOVA LNPs or AP23C5_25+ mOVA cLNPs.

**Figure 17** Gating strategy for flow cytometry plots for analysis of CD3$^{+}$CD8$^{+}$TNFα$^{+}$ cells and CD3+CD8+IFNγ+ cells in the spleen after vaccination.

**Figure 18** Representative flow cytometry plots for analysis of CD3$^{+}$ CD8$^{+}$ IFN-γ$^{+}$ cells in the spleen by the SM-102 mOVA LNPs or AP23C5_25+ mOVA cLNPs.

**Figure 19** Representative flow cytometry plots for analysis of CD3$^{+}$CD8$^{+}$TNFα$^{+}$ cells in the spleen by the SM-102 mOVA LNPs or AP23C5_25+ mOVA cLNPs.

**Figure 20** Representative flow cytometry plots for determining OVA-specific $CD8^+$ T cells in the spleen are shown by the ALC-0315 mOVA LNPs or AP23C5_25+ mOVA cLNPs.

**Figure 21** Representative flow cytometry plots for analysis of $CD3^+$ $CD8^+$ IFN-$\gamma^+$ cells in the spleen by the ALC-0315 mOVA LNPs or AP23C5_25+ mOVA cLNPs.

**Figure 22** Representative flow cytometry plots for analysis of $CD3^+CD8^+TNF\alpha^+$ cells in the spleen by the ALC-0315 mOVA LNPs or AP23C5_25+ mOVA cLNPs.

**Figure 23** Representative images of IFN-γ-secreting cells from the enzyme-linked immunosorbent assay.

**Figure 24** *In vivo* assessment of lyophilized cLNP formulations for enhanced mRNA vaccine immunity.

**Figure 25** Anti-tumour efficacy of crosslinked mRNA LNP formulations as therapeutic vaccines for B16-OVA tumor model.

**Figure 26** Anti-tumour efficacy of crosslinked mRNA LNP formulations as therapeutic vaccines for B16F10 tumor model.

**Table 1** Composition details of the evaluated LNP formulations and their corresponding crosslinked AP23C5_25+ cLNPs.

#### Synthesis of 5-hydroxymethyl-2,2,5-trimethyl-1,3-dioxane (CDCL1)

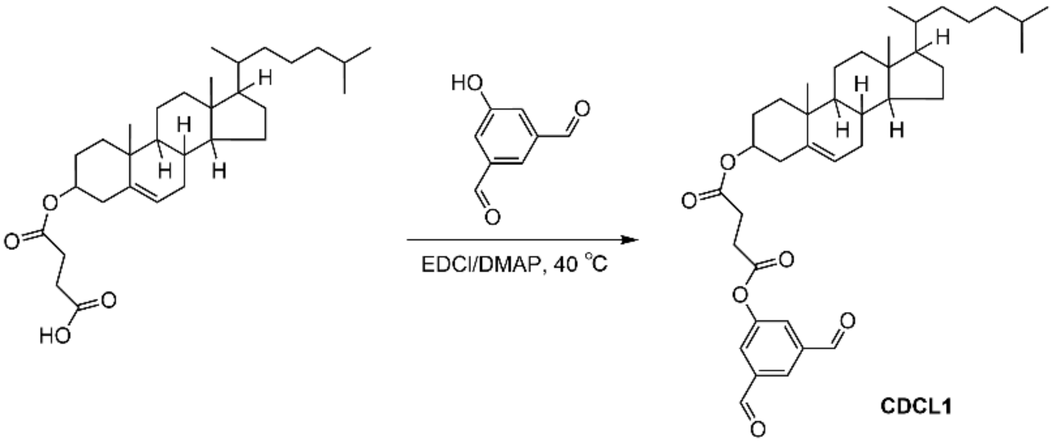


**Scheme S1.** Synthesis of compound CDCL1.

CHEMS (0.6 g, 1.23 mmol), EDC (0.555 g, 2.43 mmol), and DMAP (29.2 mg, 0.24 mmol) were dispersed in anhydrous dichloromethane with stirring for 1 h, and then 5-hydroxyisophthalaldehyde (0.234 g, 1.56 mmol) was added dropwise and reacted at 40°C overnight. The reaction solution was filtered, and all the filtrate was retained. The crude product was purified by column chromatography using dichloromethane/methanol as the eluent (0–10% methanol). The column fraction containing the purified product was separated, and the solvent was removed under reduced pressure to yield CDCL1 as a white solid. $^{1}$H-NMR spectrum of the synthesized product was obtained with a Bruker Avance III 400 MHz instrument. Characterization Data: $^{1}$H NMR (400 MHz, DMSO-d6): δ 10.12 (s, 2H), 8.38 (s, 1H), 7.96 (s, 2H), 5.35 (s, 1H), 4.52 (s, 1H), 2.80 (d, J = 79.9 Hz, 4H), 2.29 (s, 2H), 2.09 – 0.74 (m, 38H), 0.66 (s, 3H) ppm.

**Synthesis of 5-hydroxymethyl-2,2,5-trimethyl-1,3-dioxane (CDCL2)**

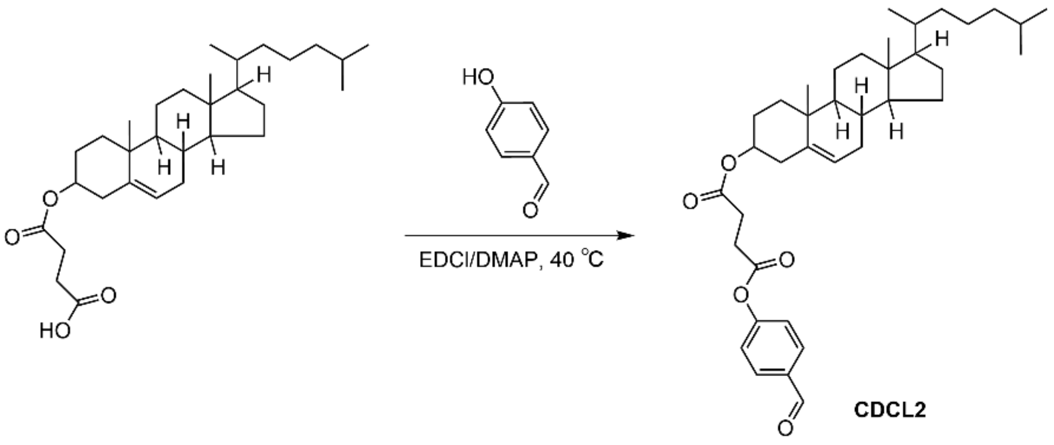


**Scheme S2.** Synthesis of compound CDCL2.

CHEMS (0.6 g, 1.23 mmol), EDC (0.555 g, 2.43 mmol), and DMAP (29.2 mg, 0.24 mmol) were dispersed in anhydrous dichloromethane with stirring for 1 h, and then 4-hydroxy-benzaldehyde (0.19 g, 1.56 mmol) was added dropwise and reacted at 40°C overnight. The reaction solution was filtered, and all the filtrate was retained. The crude product was purified by column chromatography using dichloromethane/methanol as the eluent (0–10% methanol). The column fraction containing the purified product was separated, and the solvent was removed under reduced pressure to yield CDCL2 as a white solid. $^1$H-NMR spectrum of the synthesized product was obtained with a Bruker Avance III 400 MHz instrument. Characterization Data: $^1$H NMR (400 MHz, $CDCl_3$): δ 10.00 (s, 1H), 7.92 (d, $J$ = 8.7 Hz, 2H), 7.30 (d, $J$ = 8.6 Hz, 2H), 5.38 (d, $J$ = 4.1 Hz, 1H), 4.67 (m, 1H), 3.02 – 2.61 (m, 4H), 2.34 (d, $J$ = 8.0 Hz, 2H), 2.08 – 0.78 (m, 38H), 0.68 (s, 3H) ppm.

#### Synthesis of CDCL3

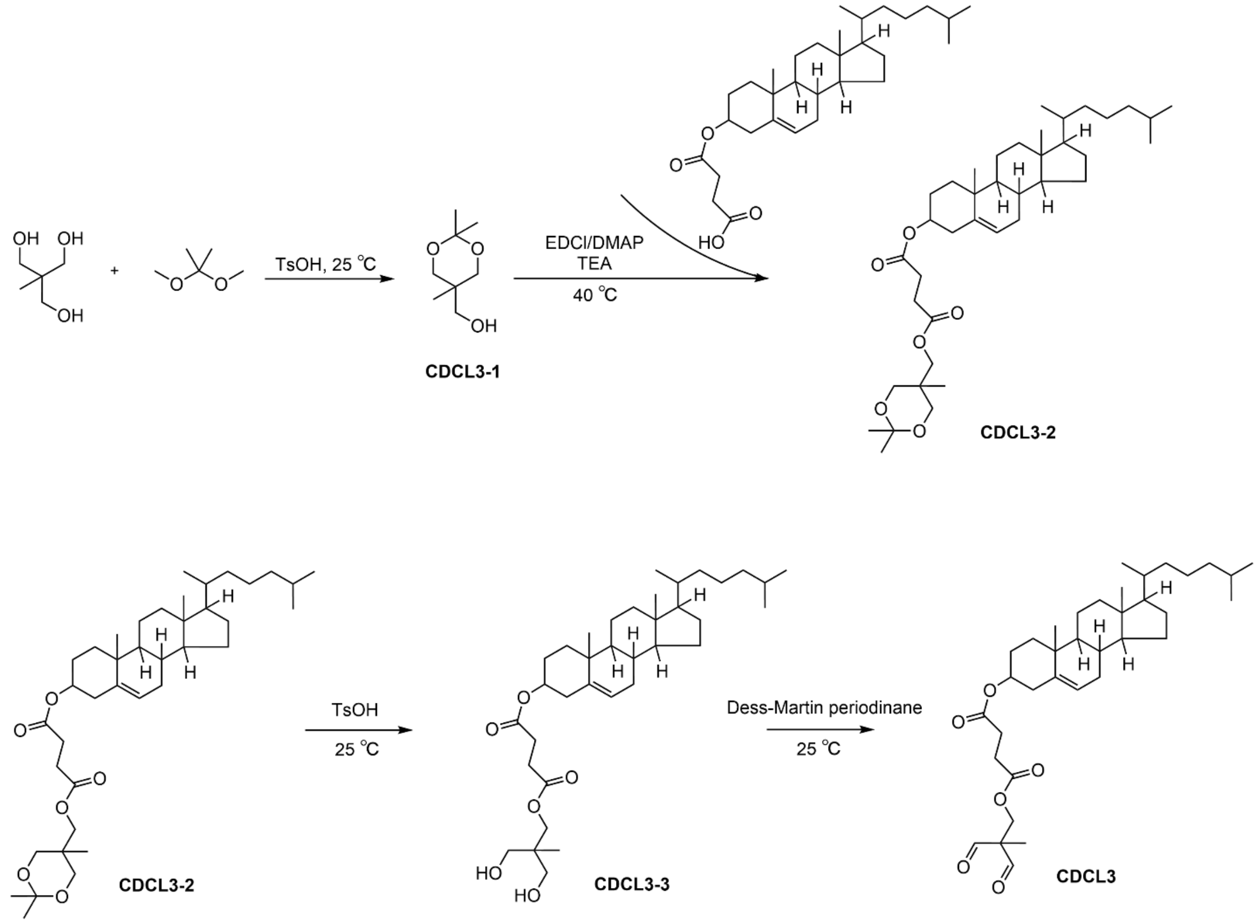


**Scheme S3.** Synthesis of compound CDCL3.

1,1,1-tris(hydroxymethyl)ethane (10 g, 83.22 mmol), 2,2-dimethoxypropane (8.66 g, 83.22 mmol), p-TsOH (158 mg, 0.8 mmol), and acetone (80 mL) to a 250 mL round bottom flask and stir for 5 h at room temperature. After that, $K_2CO_3$ solution was added to the reaction solution. The reaction solution was filtered, and all the filtrate was retained. The crude product was then purified by column chromatography using n-hexane/EtOAc as the eluent (0–20% EtOAc). The column fraction containing the purified product was separated, and the solvent was removed under reduced pressure to yield 5-hydroxymethyl-2,2,5-trimethyl-1,3-dioxane (CDCL3-1) as an oily transparent liquid. $^1$H NMR spectrum of the synthesized product was obtained with a Bruker Avance III 400 MHz instrument. Characterization Data: $^1$H NMR (400 MHz, $CDCl_3$): δ 3.83 – 3.49 (m, 6H), 1.80 (s, 1H), 1.41 (d, $J$ = 18.7 Hz, 6H), 0.82 (s, 3H) ppm.

CHEMS (0.3 g, 0.62 mmol), EDC (0.278 g, 1.22 mmol), TEA (0.147 g, 1.45 mmol) and DMAP (14.6 mg, 0.12 mmol) were dispersed in anhydrous dichloromethane with stirring for 1 h, and then CDCL3-1 (0.125 g, 0.78 mmol) was added dropwise and reacted at 40°C overnight. The reaction solution was filtered, and all the filtrate was retained. The crude product was purified by column chromatography using dichloromethane/methanol as the eluent (0–10% methanol). The column fraction containing the purified product was separated, and the solvent was removed under reduced pressure to yield CDCL3-2 as a white solid. $^1$H NMR spectrum of the synthesized product was obtained with a Bruker Avance III 400 MHz instrument. Characterization Data: $^{1}$H NMR (400 MHz, $CDCl_3$): δ 5.37 (d, *J* = 4.1 Hz, 1H), 4.62 (m, 1H), 4.19 (s, 2H), 3.63 (q, *J* = 11.9 Hz, 4H), 2.63 (dd, *J* = 11.8, 6.2 Hz, 4H), 2.31 (d, *J* = 7.8 Hz, 2H), 2.08 – 0.78 (m, 47H), 0.67 (s, 3H) ppm.

CDCL3-2 (520 mg, 0.83 mmol) and p-TsOH (17.3 mg, 0.1 mmol) were dispersed in anhydrous dichloromethane with stirring for 6 h at room temperature, and then the reaction solution was filtered, and all the filtrate was retained. The crude product was purified by column chromatography using dichloromethane/methanol as the eluent (0–10% methanol). The column fraction containing the purified product was separated, and the solvent was removed under reduced pressure to yield CDCL3-3 as a transparent solid. $^{1}$H NMR spectrum of the synthesized product was obtained with a Bruker Avance III 400 MHz instrument. Characterization Data: $^{1}$H NMR (400 MHz, $CDCl_3$): δ 5.37 (d, *J* = 3.4 Hz, 1H), 4.62 (m, 1H), 4.23 (s, 2H), 3.55 (q, *J* = 11.2 Hz, 4H), 2.76 (s, 2H), 2.69 – 2.56 (m, 4H), 2.31 (d, *J* = 7.5 Hz, 2H), 2.08 – 0.78 (m, 41H), 0.67 (s, 3H) ppm.

CDCL3-3 (180 mg, 0.31 mmol) and DMP (387.47 mg, 0.91 mmol) were dispersed in anhydrous dichloromethane with stirring for 2 h at room temperature. After that, saturated sodium bicarbonate and saturated sodium bisulfite solutions were added to quench the reaction. The precipitate was then filtered, and the filtrate was layered to extract the oil phase. The crude product was purified by column chromatography using dichloromethane/methanol as the eluent (0–5% methanol). The column fraction containing the purified product was separated, and the solvent was removed under reduced pressure to yield CDCL3 as a white solid. $^{1}$H NMR spectrum of the synthesized product was obtained with a Bruker Avance III 400 MHz instrument. Characterization Data: $^{1}$H NMR (400 MHz, $CDCl_3$): δ 9.70 (s, 2H), 5.36 (s, 1H), 4.62 (m, 1H), 2.59 (m, 4H), 2.30 (d, *J* = 7.6 Hz, 2H), 2.20 – 0.77 (m, 41H), 0.67 (s, 3H) ppm.

**Synthesis of CDCL4**

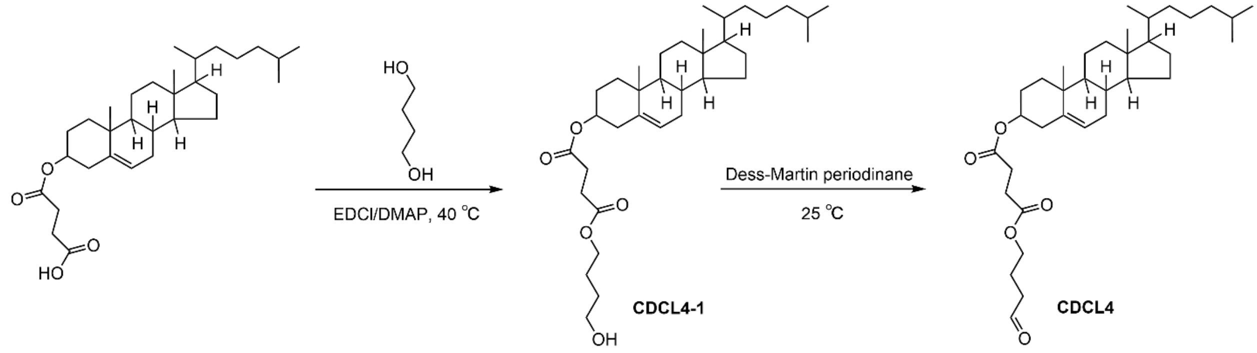


**Scheme S4.** Synthesis of compound CDCL4.

CHEMS (0.5 g, 1.03 mmol), EDC (0.463 g, 2.03 mmol) and DMAP (24.3 mg, 0.2 mmol) were dispersed in anhydrous dichloromethane with stirring for 1 h, and then butane-1,4-diol (92.4 mg, 1.03 mmol) was added dropwise and reacted at 40°C overnight. The reaction solution was filtered, and all the filtrate was retained. The crude product was purified by column chromatography using dichloromethane/methanol as the eluent (0-10% methanol). The column fraction containing the purified product was separated, and the solvent was removed under reduced pressure to yield CDCL4-1 as a white solid. $^1$H-NMR spectrum of the synthesized product was obtained with a Bruker Avance III 400 MHz instrument. Characterization Data: $^1$H NMR (400 MHz, $CDCl_3$): $^1$H NMR (400 MHz, $CDCl_3$): δ 5.37 (s, 1H), 4.63 (m, 1H), 4.26 – 3.95 (m, 2H), 3.68 (q, $J$ = 6.0 Hz, 2H), 2.61 (d, $J$ = 1.4 Hz, 4H), 2.31 (d, $J$ = 7.2 Hz, 2H), 2.11 – 0.77 (m, 42H), 0.67 (s, 3H) ppm.

CDCL4-1 (196 mg, 0.35 mmol) and DMP (267.8 mg, 0.63 mmol) were dispersed in anhydrous dichloromethane with stirring for 2 h at room temperature. After that, saturated sodium bicarbonate and saturated sodium bisulfite solutions were added to quench the reaction. The precipitate was then filtered, and the filtrate was layered to extract the oil phase. The crude product was purified by column chromatography using dichloromethane/methanol as the eluent (0-5% methanol). The column fraction containing the purified product was separated, and the solvent was removed under reduced pressure to yield CDCL4 as a white solid. $^1$H-NMR spectrum of the synthesized product was obtained with a Bruker Avance III 400 MHz instrument. Characterization Data: $^1$H NMR (400 MHz, $CDCl_3$): 9.78 (s, 1H), δ 5.36 (d, $J$ = 5.1 Hz, 1H), 4.60 (m, 1H), 4.13 (d, $J$ = 12.7 Hz, 2H), 2.61 (d, $J$ = 2.9 Hz, 4H), 2.31 (d, $J$ = 7.1 Hz, 2H), 2.12 – 0.76 (m, 40H), 0.67 (s, 3H) ppm.

**Synthesis of CDCL5**

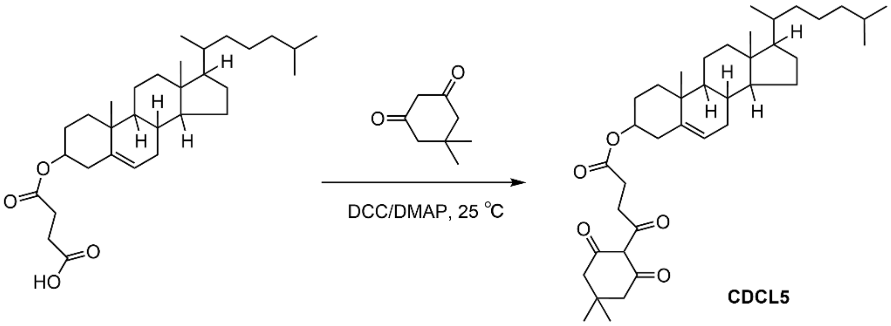


CDCL5

**Scheme S5.** Synthesis of compound CDCL5.

Briefly, 5,5-Dimethyl-1,3-cyclohexanedione (95 mg, 0.68 mmol), CHEMS (0.3 g, 0.62 mol), and DMAP (0.113 g, 0.92 mol) were dissolved in dichloromethane with stirring. A separate solution of DCC (0.152 g, 0.74 mol) in dichloromethane was added slowly at room temperature to the reaction mixture. The reaction proceeded for 4 h with stirring at room temperature, at which point the white N,N'-dicyclohexylurea precipitate was filtered off, and the precipitate was washed with dichloromethane until colorless. The dichloromethane filtrate was collected and washed with 3% HCl until the pH of the aqueous phase was < 3. The organic phase was separated, dried over MgSO4, and filtered, and the solvent was then removed under vacuum. The crude product was purified by column chromatography using dichloromethane/methanol as the eluent (0–10% methanol). The column fraction containing the purified product was separated, and the solvent was removed under reduced pressure to yield CDCL5 as a white solid. $^{1}$H NMR spectrum of the synthesized product was obtained with a Bruker Avance III 400 MHz instrument. Characterization Data: $^{1}$H NMR (400 MHz, $CDCl_3$): δ 5.36 (d, $J$ = 5.0 Hz, 1H), 4.61 (m, 1H), 3.44 – 2.56 (m, 4H), 2.44 (d, $J$ = 72.1 Hz, 4H), 2.31 (d, $J$ = 7.4 Hz, 2H), 2.09 – 0.79 (m, 44H), 0.67 (s, 3H) ppm.

#### Synthesis of Cholesteryl Hemisuccinate-N-hydroxysuccinimide (CHNH)

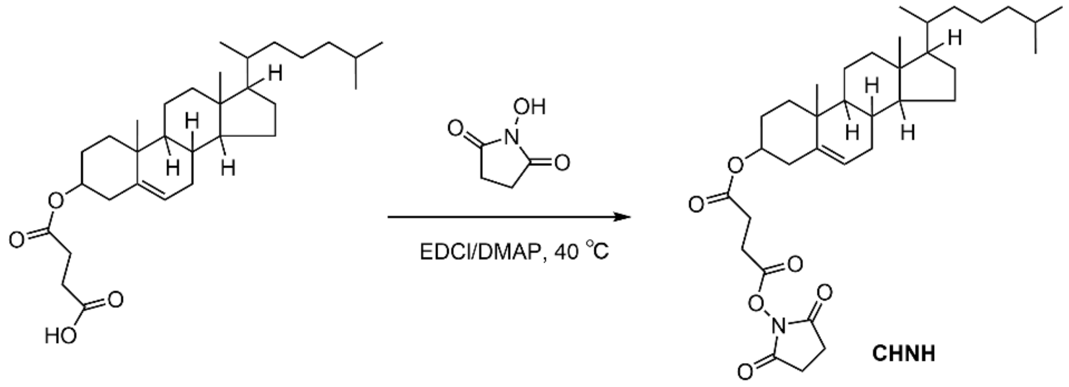


**Scheme S6.** Synthesis of compound CHNH.

CHEMS (0.6 g, 1.23 mmol), EDC (0.555 g, 2.43 mmol), and DMAP (29.2 mg, 0.24 mmol) were dispersed in anhydrous dichloromethane with stirring for 1 h, and then N-hydroxysuccinimide (0.179 g, 1.56 mmol) was added dropwise and reacted at 40°C overnight. The reaction solution was filtered, and all the filtrate was retained. The crude product was purified by column chromatography using dichloromethane/methanol as the eluent (0–10% methanol). The column fraction containing the purified product was separated, and the solvent was removed under reduced pressure to yield CHNH as a white solid. $^{1}$H-NMR spectrum of the synthesized product was obtained with a Bruker Avance III 400 MHz instrument. Characterization Data: $^{1}$H NMR (300 MHz, $CDCl_3$): δ 5.37 (d, $J$ = 5.5 Hz, 1H), 4.66 (m, 1H), 2.94 (t, $J$ = 6.8 Hz, 2H), 2.83 (m, 4H), 2.71 (t, $J$ = 7.3 Hz, 2H), 2.32 (d, $J$ = 8.0 Hz, 2H), 2.14 – 0.79 (m, 38H), 0.67 (s, 3H) ppm.

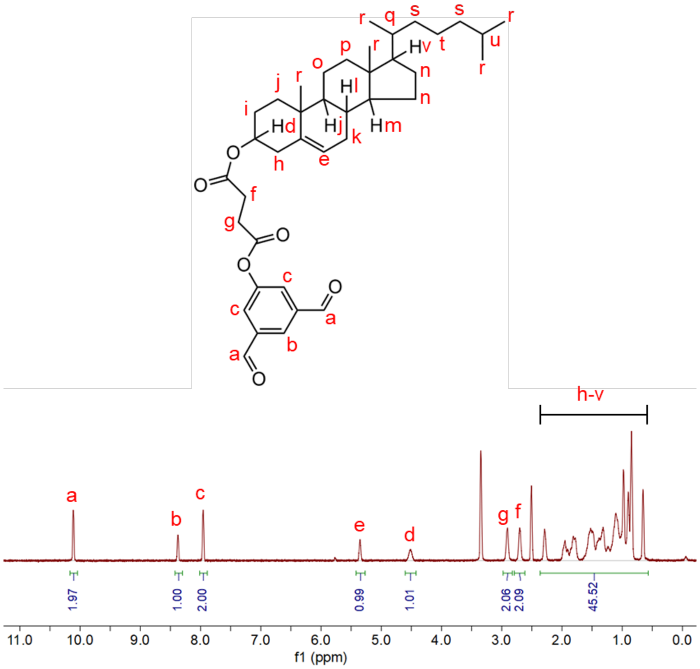


**Supplementary Figure S1.** Representative $^{1}$H-NMR spectrum of CDCL1.

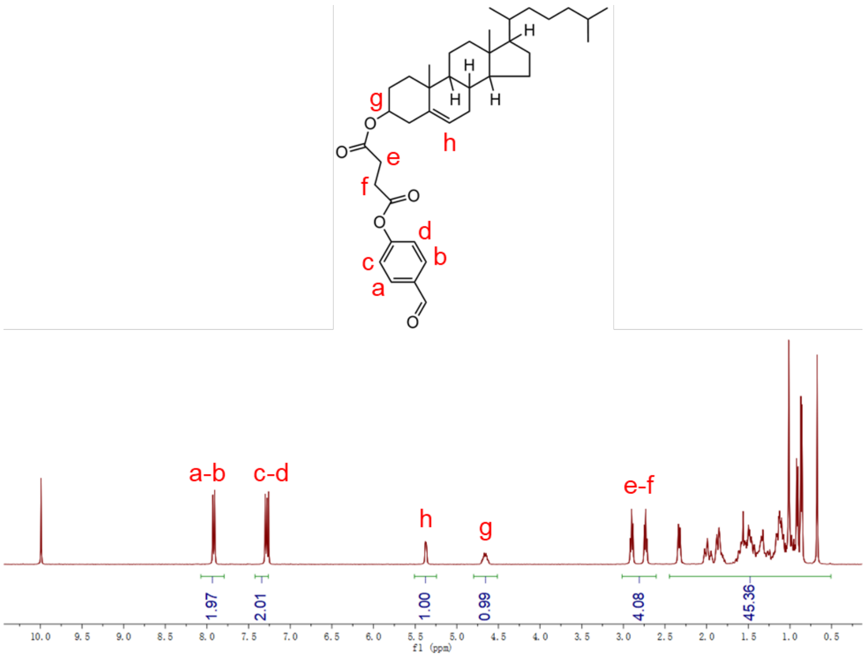


**Supplementary Figure S2.** Representative $^1$H-NMR spectrum of CDCL2.

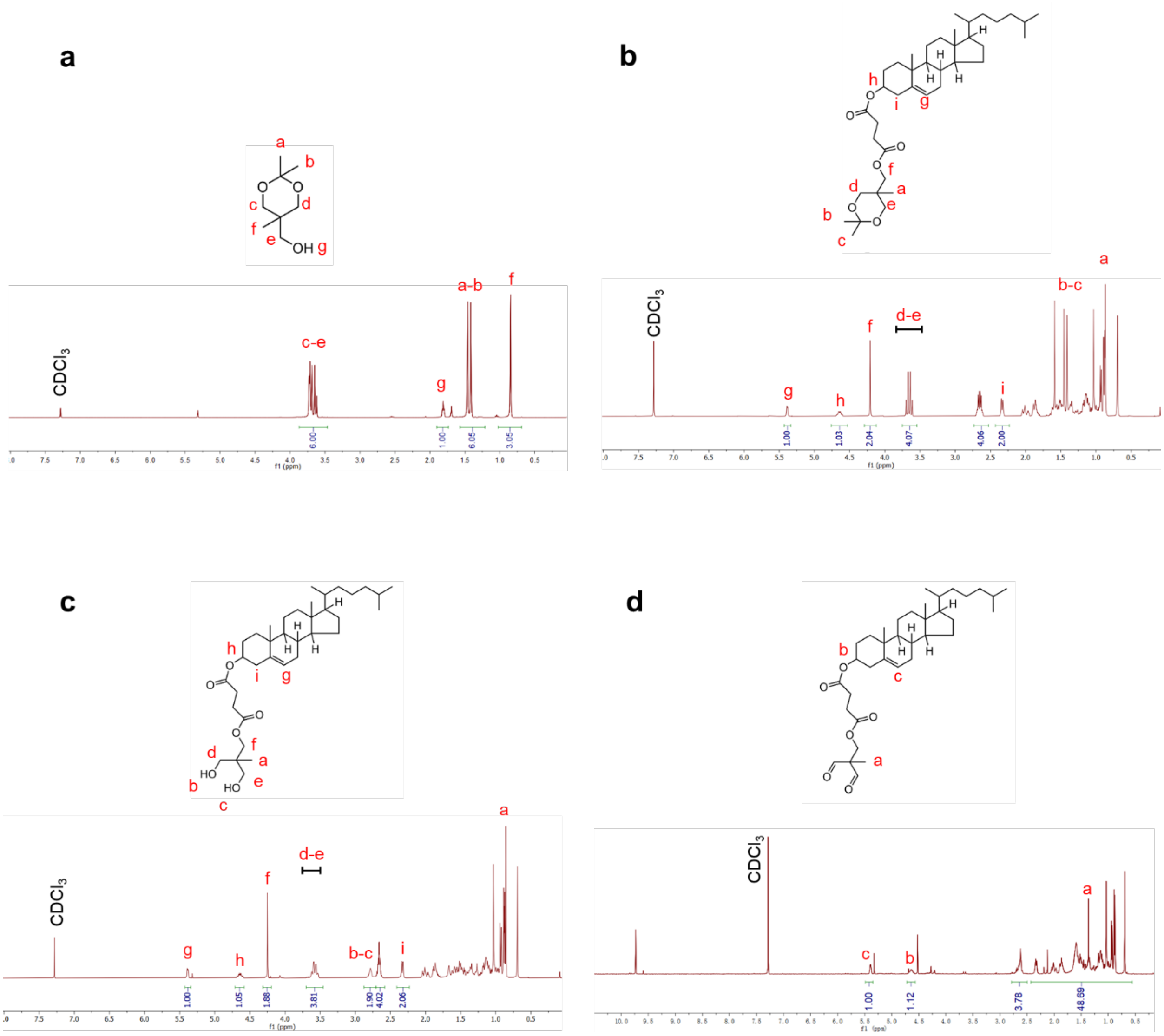


**Supplementary Figure S3. a.** Representative $^{1}$H-NMR spectrum of CDCL3-1. **b.** Representative $^{1}$H-NMR spectrum of CDCL3-2. **c.** Representative $^{1}$H-NMR spectrum of CDCL3-3. **d.** Representative $^{1}$H-NMR spectrum of CDCL3.

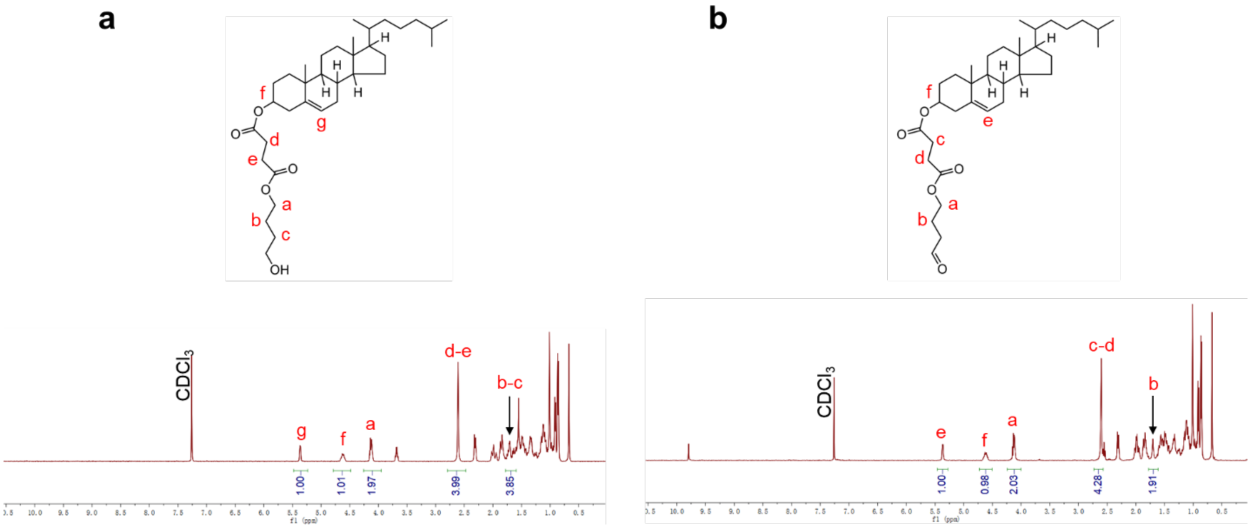


**Supplementary Figure S4. a.** Representative $^1$H-NMR spectrum of CDCL4-1. **b.** Representative $^1$H-NMR spectrum of CDCL4.

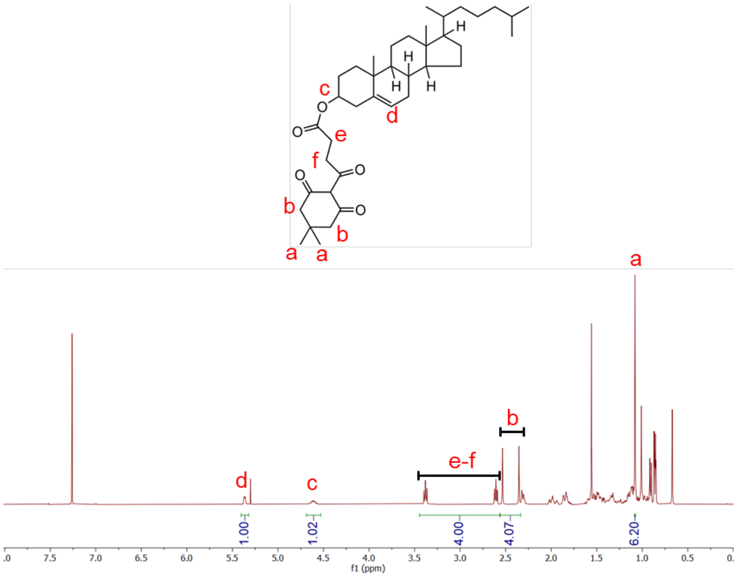


**Supplementary Figure S5.** Representative [1]H-NMR spectrum of CDCL5.

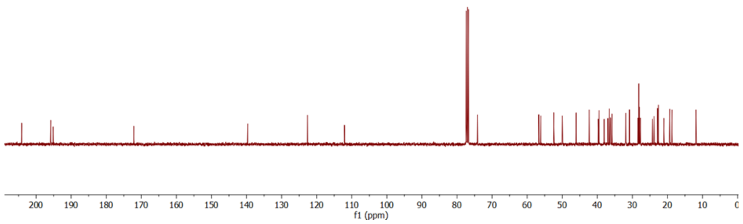


**Supplementary Figure S6.** Representative $^{13}C$ NMR spectrum of CDCL5.

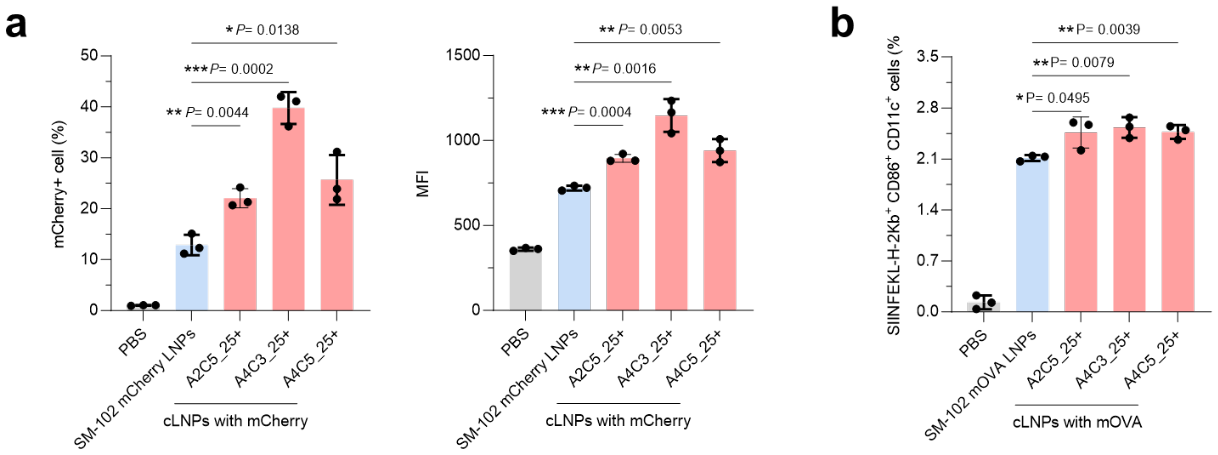


**Supplementary Figure S7.** ***In vitro*** **evaluation of top-performing crosslinking candidates in SM-102 LNPs for transfection and maturation of DCs. a.** BMDCs were treated with top-performing crosslinked LNPs or original SM-102 LNP formulations loaded with mCherry mRNA. The figure illustrates the percentage and the mean fluorescence intensity (MFI) of mCherry-positive cells after 24 h of incubation with mRNA cLNPs. **b.** The maturation of BMDCs was evaluated using flow cytometry following 24 h incubation with three top-performing cLNP or original SM-102 LNP formulations loaded with mOVA.

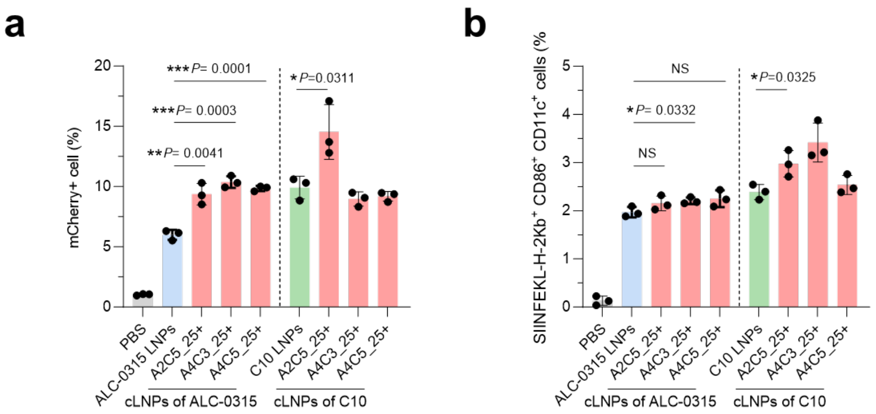


**Supplementary Figure S8. *In vitro* evaluation of top-performing crosslinking candidates in ALC-0315 LNPs or C10 LNPs for transfection and maturation of DCs. a.** BMDCs were treated with top-performing cLNPs or original LNP formulations loaded with mCherry mRNA. The figure illustrates the percentage of mCherry-positive cells after 24 h of incubation with mRNA cLNPs. **b.** The maturation of BMDCs was evaluated using flow cytometry following 24 h incubation with three top-performing cLNP or original LNP formulations loaded with mOVA.

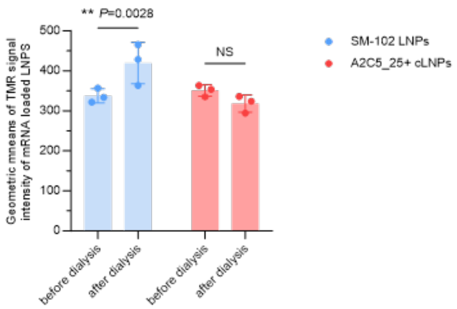


**Supplementary Figure S9.** The geometric means of TMR signal (indicator of relative helper lipid content) intensity of mRNA loaded LNPs.

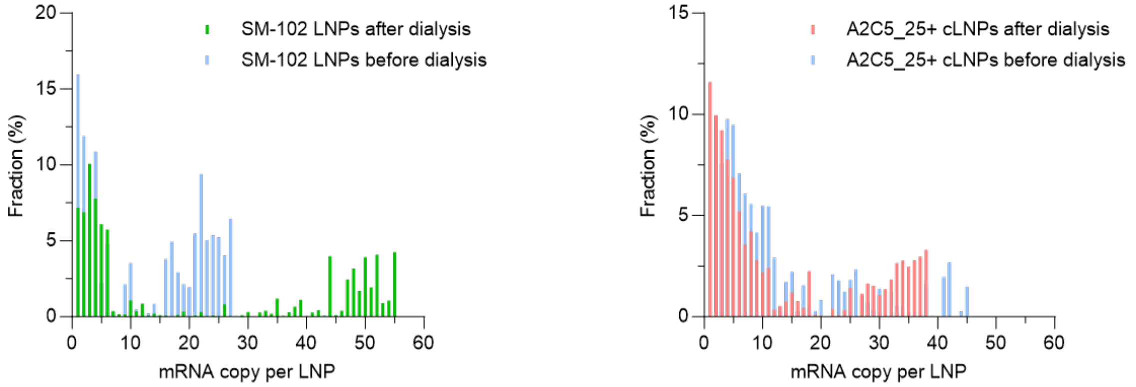


**Supplementary Figure S10.** The mRNA payload distribution profiles of SM-102 LNP or A2C5_25+ cLNP formulations before and after dialysis.

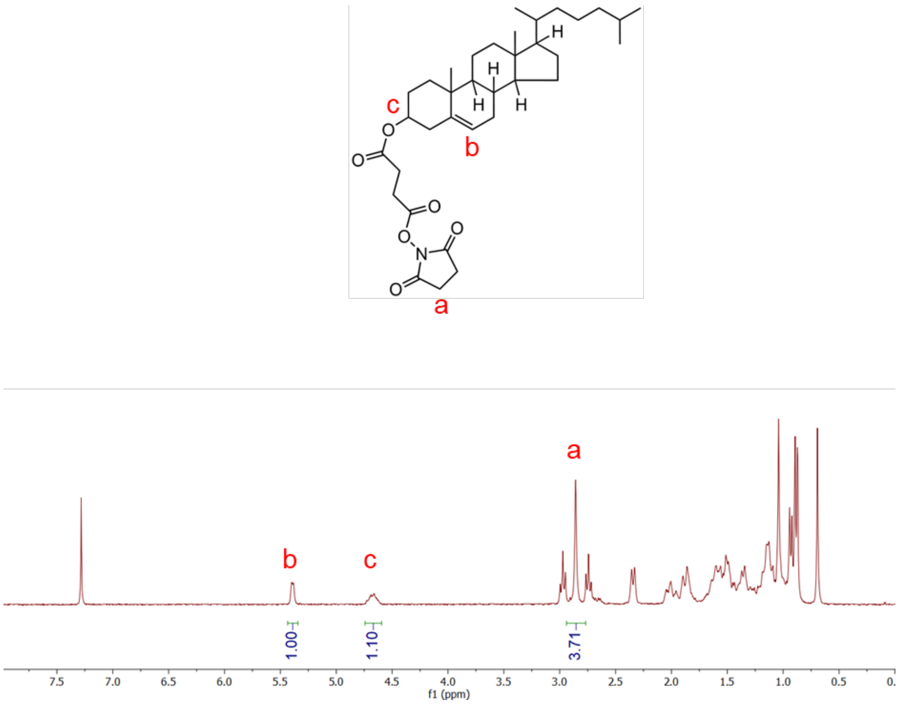


**Supplementary Figure S11.** Representative $^{1}$H-NMR spectrum of CHNH.

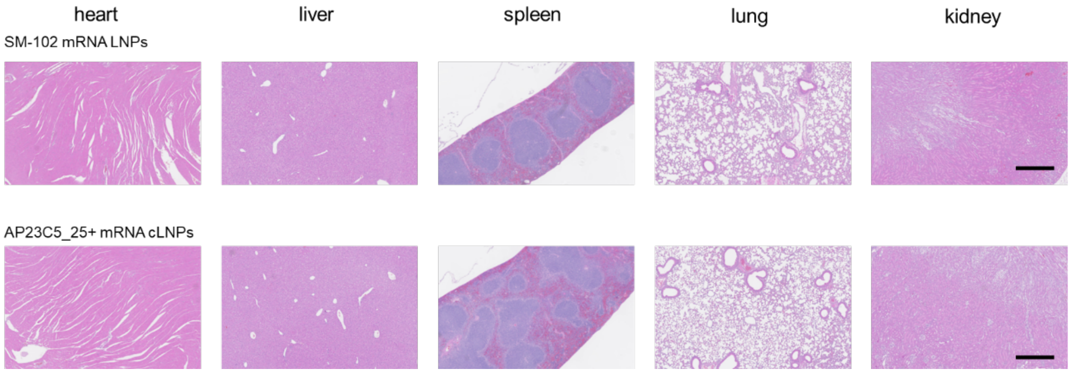


**Supplementary Figure S12.** H&E staining of major organs of mice treated with SM-102 LNPs or AP23C5_25+ LNPs for in vivo biosafety. Scale bar, 500 µm.

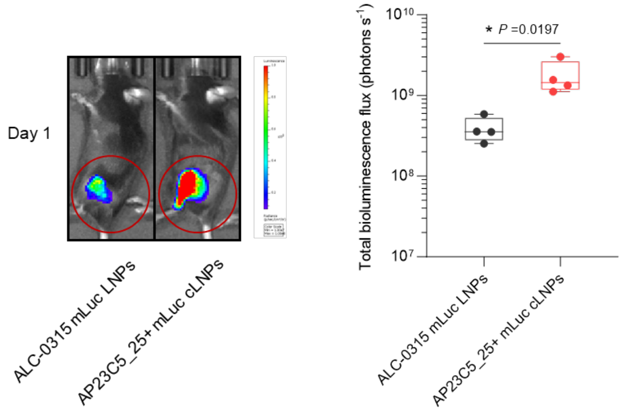


**Supplementary Figure S13. *In vivo* transfection efficiency of crosslinked vs. uncrosslinked ALC-0315 LNPs.** ALC-0315 LNP and AP23C5_25+ cLNP formulations were *i.m.* injected into mice, with luciferase expression at the injection site imaged and quantified for total luminescent flux on Day 1 post-injection using the IVIS (n = 4 biologically independent samples). A representative bioluminescence image from Day 1 is presented. For boxplots, the box extends from the 25th to the 75th percentiles, and the line in the middle of the box is plotted at the median. Data were analyzed using a two-tailed Student's t-test between two. *$P < 0.05$, **$P < 0.01$, ***$P < 0.001$, ****$P < 0.0001$.

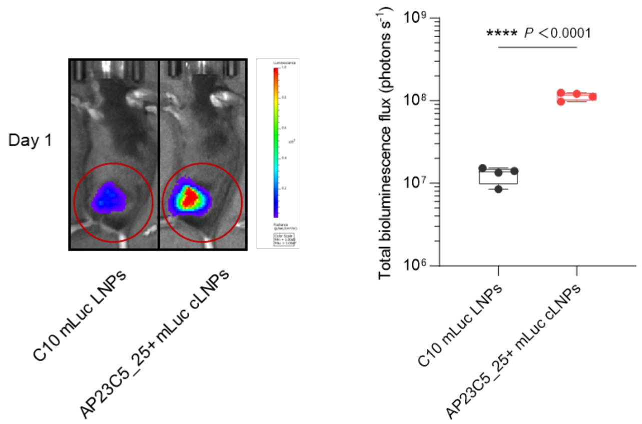


**Supplementary Figure S14. *In vivo* transfection efficiency of crosslinked vs. uncrosslinked C10 LNPs.** C10 LNP and AP23C5_25+ cLNP formulations were *i.m.* injected into mice, with luciferase expression at the injection site imaged and quantified for total luminescent flux on Day 1 post-injection using the IVIS (n = 4 biologically independent samples). A representative bioluminescence image from Day 1 is presented. For boxplots, the box extends from the 25th to the 75th percentiles, and the line in the middle of the box is plotted at the median. Data were analyzed using a two-tailed Student's t-test between two. $*P < 0.05$, $**P < 0.01$, $***P < 0.001$, $****P < 0.0001$.

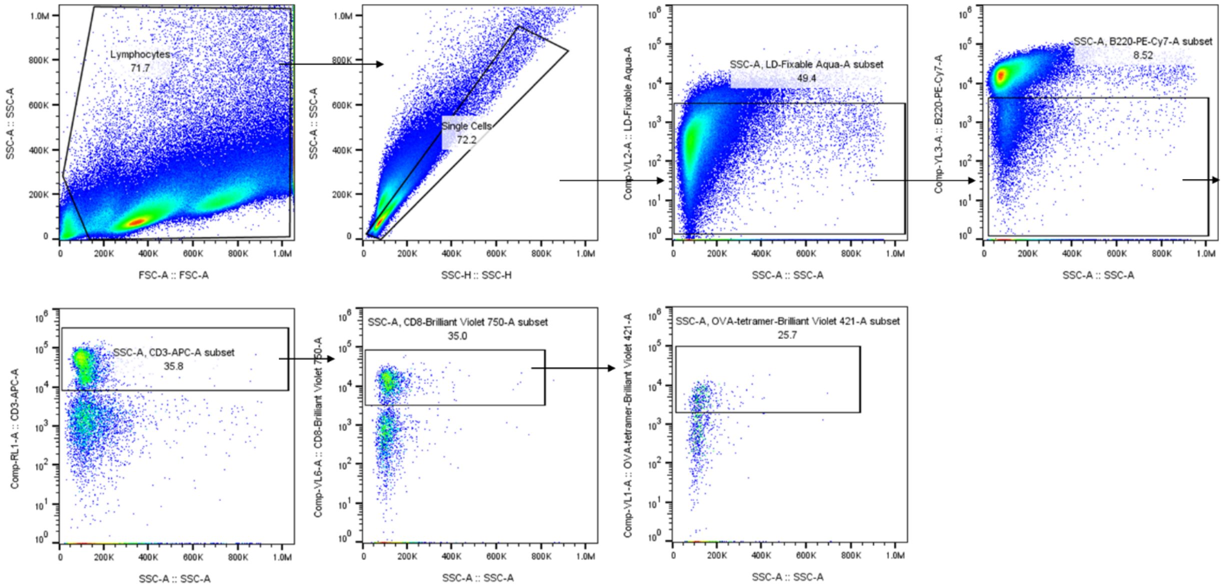


**Supplementary Figure S15.** Gating strategy for flow cytometry plots for antigen (OVA)-specific T cell responses in the spleen after vaccination.

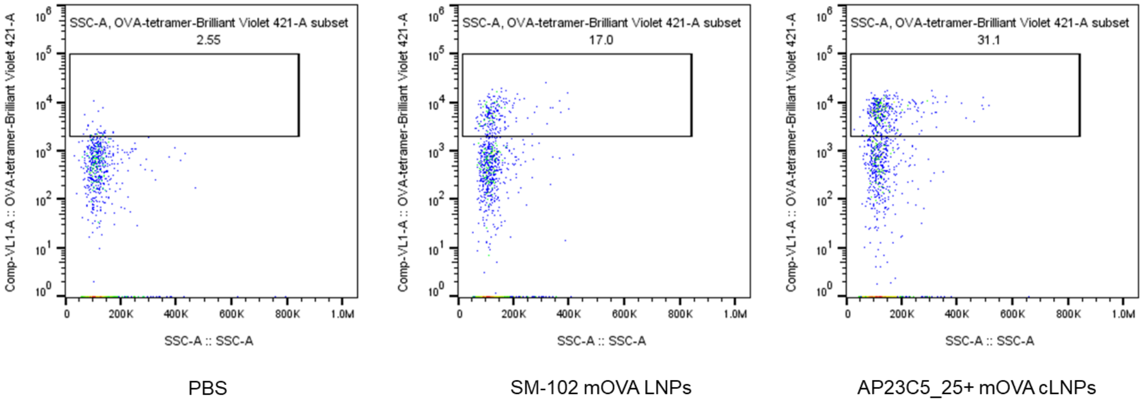


**Supplementary Figure S16.** Representative flow cytometry plots for determining OVA-specific CD8$^{+}$ T cells in the spleen are shown by the SM-102 mOVA LNPs or AP23C5_25+ mOVA cLNPs.

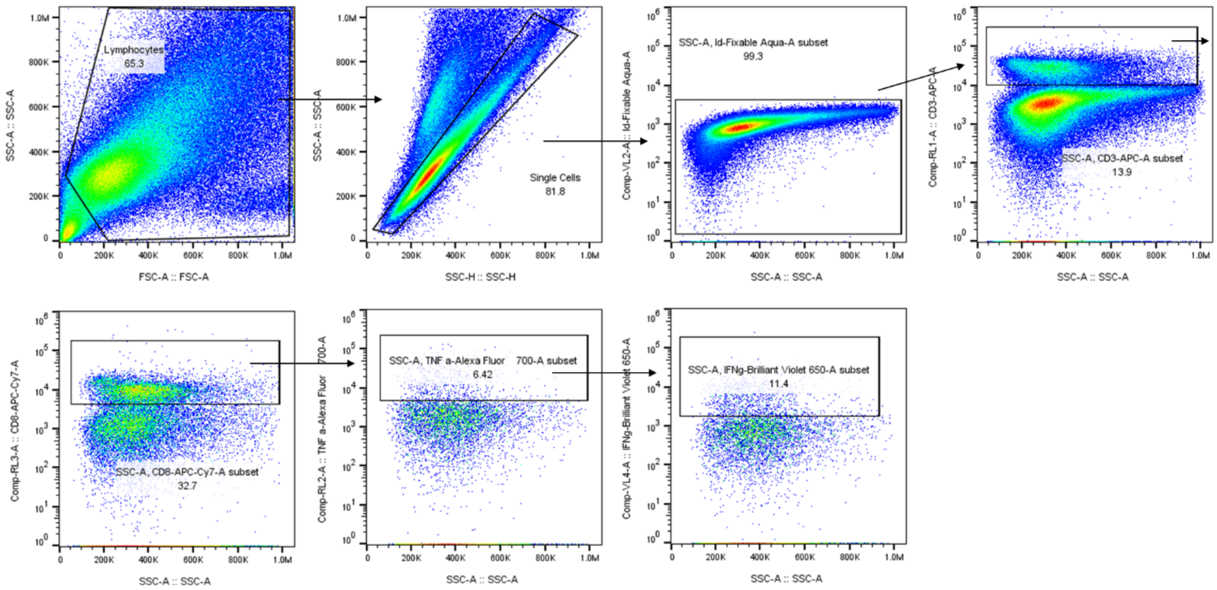


**Supplementary Figure S17.** Gating strategy for flow cytometry plots for analysis of $CD3^{+}CD8^{+}TNF\alpha^{+}$ cells and $CD3^{+}CD8^{+}IFN\gamma^{+}$ cells in the spleen after vaccination.

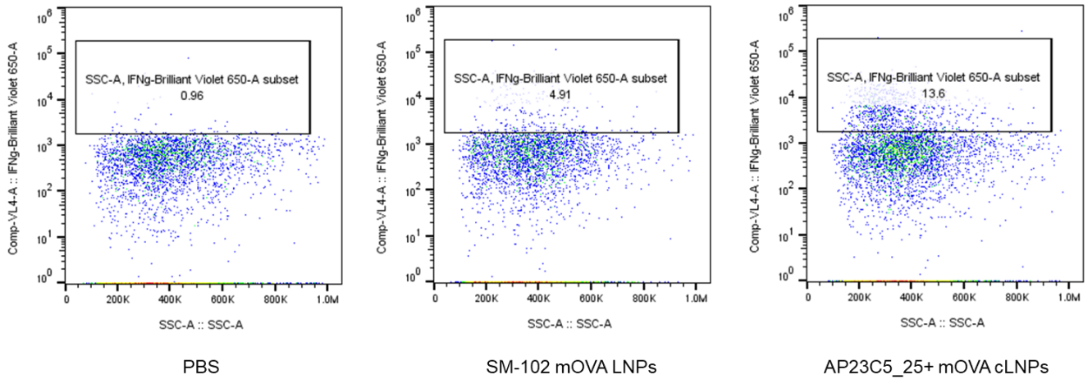


**Supplementary Figure S18.** Representative flow cytometry plots for analysis of CD3$^+$ CD8$^+$ IFN-$\gamma^+$ cells in the spleen by the SM-102 mOVA LNPs or AP23C5_25+ mOVA cLNPs.

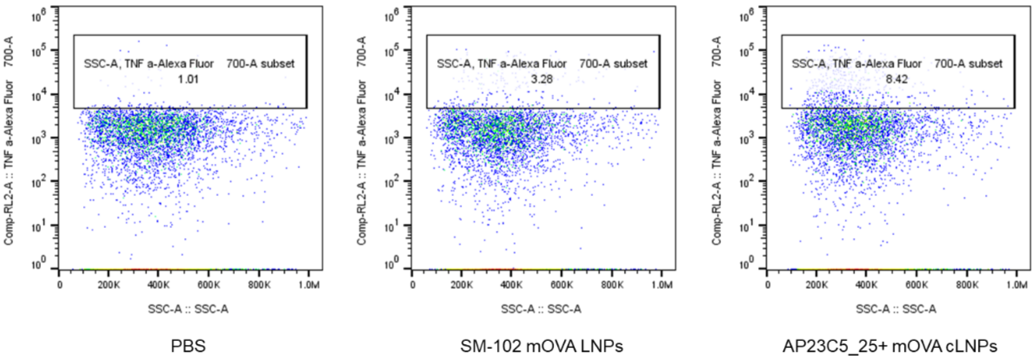


**Supplementary Figure S19.** Representative flow cytometry plots for analysis of $CD3^+CD8^+TNF\alpha^+$ cells in the spleen by the SM-102 mOVA LNPs or AP23C5_25+ mOVA cLNPs.

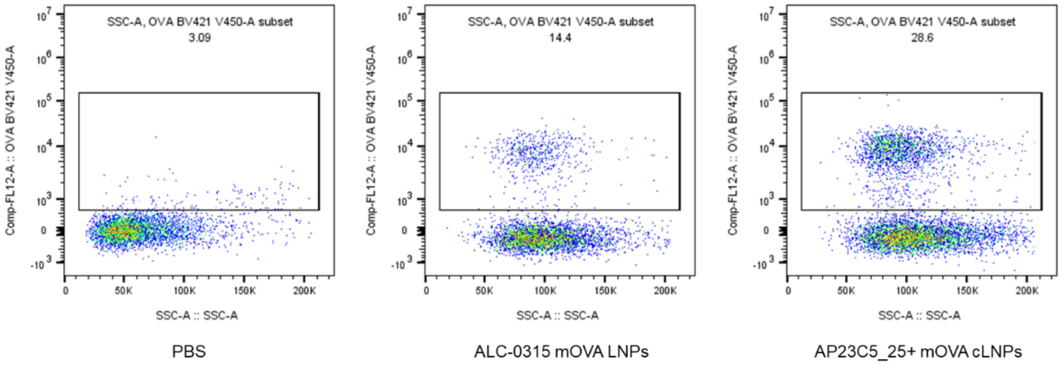


**Supplementary Figure S20.** Representative flow cytometry plots for determining OVA-specific $CD8^+$ T cells in the spleen are shown by the ALC-0315 mOVA LNPs or AP23C5_25+ mOVA cLNPs.

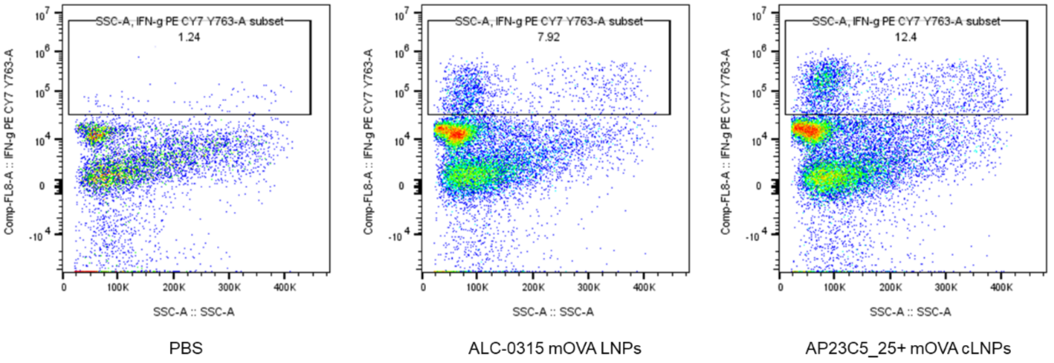


**Supplementary Figure S21.** Representative flow cytometry plots for analysis of $CD3^{+}$ $CD8^{+}$ IFN-$\gamma^{+}$ cells in the spleen by the ALC-0315 mOVA LNPs or AP23C5_25+ mOVA cLNPs.

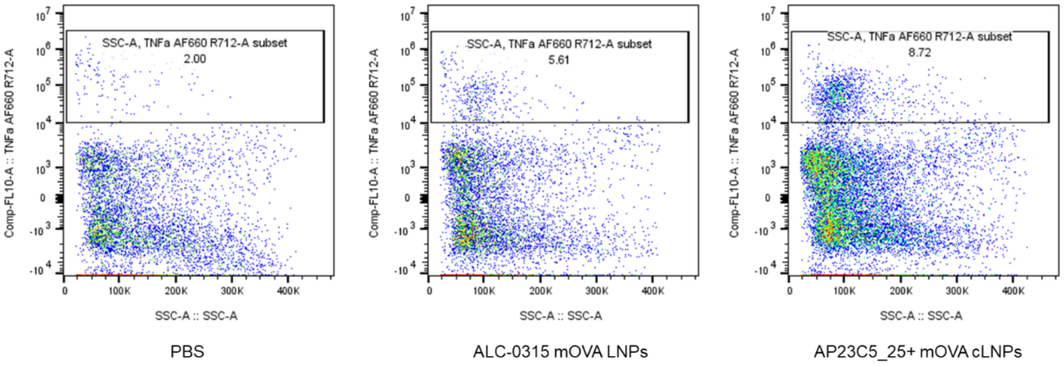


**Supplementary Figure S22.** Representative flow cytometry plots for analysis of $CD3^{+}CD8^{+}TNF\alpha^{+}$ cells in the spleen by the ALC-0315 mOVA LNPs or AP23C5_25+ mOVA cLNPs.

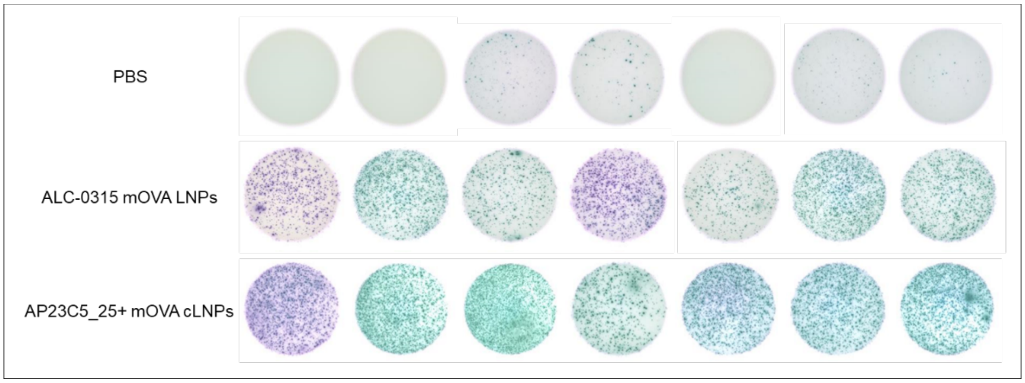


**Supplementary Figure S23. Representative images of IFN-γ-secreting cells from the enzyme-linked immunosorbent assay.** Frequency of IFN-γ-secreting cells among restimulated splenocytes, assessed via ELISpot. Splenocytes were restimulated in vitro with SIINFEKL peptide (2 µg/mL SIINFEKL).

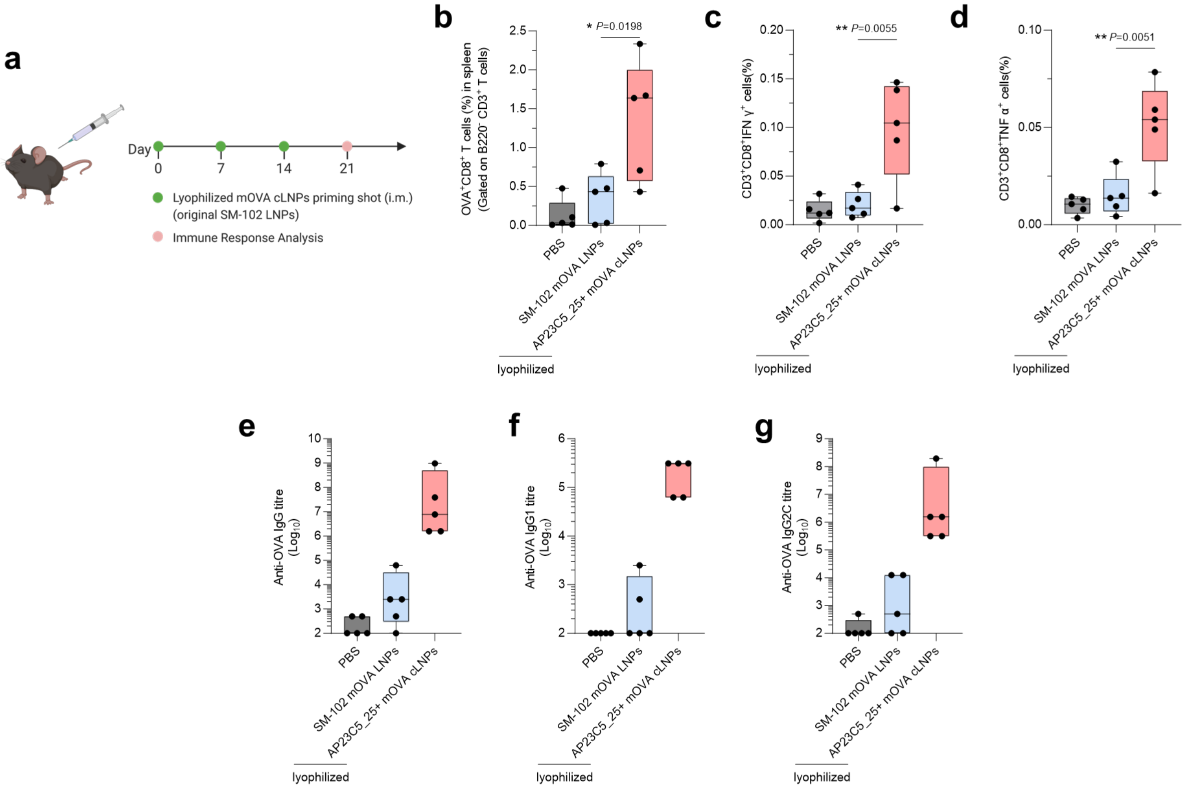


**Supplementary Figure S24.** ***In vivo*** **assessment of lyophilized cLNP formulations for enhanced mRNA vaccine immunity. a–d.** C57BL/6 mice were administered with PBS, lyophilized SM-102 LNPs, or lyophilized AP23C5_25+ cLNPs loaded with mOVA via *i.m.* injection (10 µg mOVA per injection). Mice were sacrificed 7 days after the final injection, and their splenocytes were isolated for analysis **(a)**. The percentages of OVA-specific CD8 T cells ($B220^{-}CD3^{+}CD8^{+}OVA^{+}$ cells) **(b)**. Splenocytes were restimulated *in vitro* with OVA and SIINFEKL peptide (100 µg $mL^{-1}$ OVA and 2 µg $mL^{-1}$ SIINFEKL) for 6 h and assessed via flow cytometry and intracellular cytokine staining to determine the percentages of $CD3^{+}CD8^{+}IFN\text{-}\gamma^{+}$ **(c)** and $CD3^{+}CD8^{+}TNF\text{-}\alpha^{+}$ **(d)**. **e–g.** Titers of OVA-specific IgG **(e)**, IgG1 **(f)**, and IgG2c **(g)** antibodies in blood serum on Day 21, determined by ELISA.

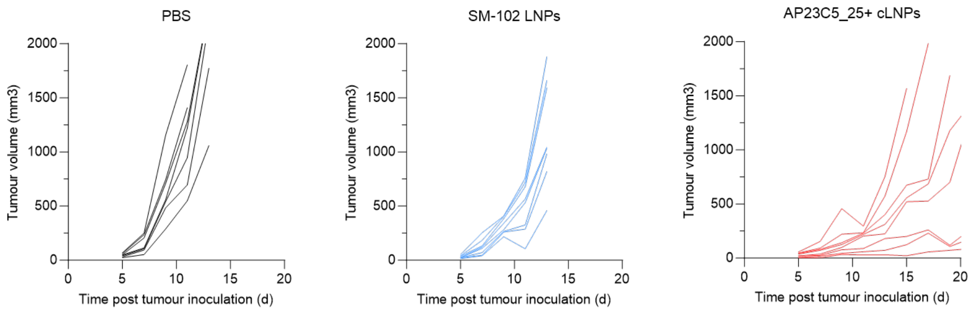


**Supplementary Figure S25. Anti-tumour efficacy of crosslinked mRNA LNP formulations as therapeutic vaccines for B16-OVA tumor model.** Mice were s.c. inoculated with B16-OVA cells and subsequently received three s.c. injections, one week apart, of PBS, mOVA-loaded SM-102 LNPs, or AP23C5_25+ SM-102 cLNPs. The results presented individual tumor volumes over time.

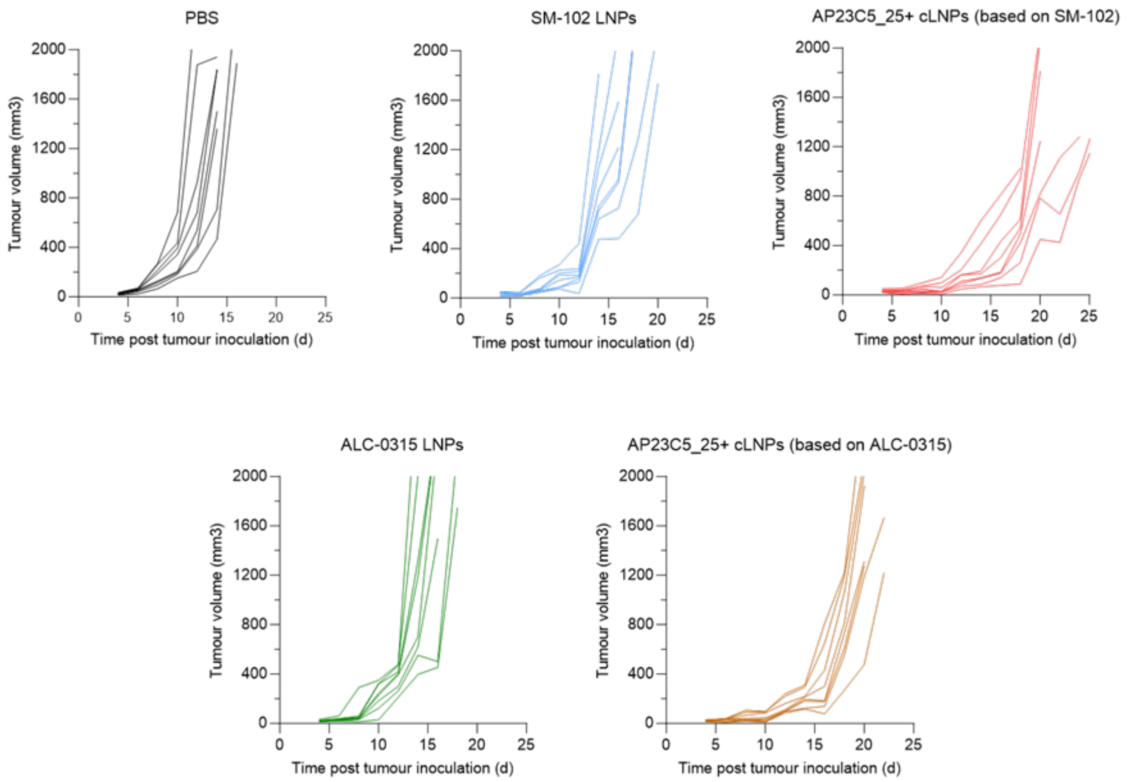


**Supplementary Figure S26. Anti-tumour efficacy of crosslinked mRNA LNP formulations as therapeutic vaccines for B16F10 tumor model.** Mice were inoculated s.c. with B16F10 cells and then given three i.m. injections, one week apart, of PBS or SM-102 LNPs, AP23C5_25+ SM-102 cLNPs, ALC-0315 LNPs, or AP23C5_25+ ALC-0315 cLNPs loaded with mRNA encoding Trp2 (mTrp) or Gp100 (mGp100) (10 µg mRNA per injection). The results presented individual tumor volumes over time.

**Supplementary Table S1. Composition details of the evaluated LNP formulations and their corresponding crosslinked AP23C5_25+ cLNPs.**

| **LNP Formula** | **SM-102 LNPs** | **AP23C5_25+ cLNPs** |
|---|---|---|
| lipid phase | SM-102:Chol:DSPC:DMG-PEG = (50 : 38.5 : 10 : 1.5) | SM-102:Chol:(CDCL5:AP23):DSPC:DMG-PEG = (50 : 38.5 : (9.63 : 9.63) : 10 : 1.5) |
| **LNP Formula** | **ALC-0315 LNPs** | **AP23C5_25+ cLNPs** |
| lipid phase | ALC-0315:Chol:DSPC:DMG-PEG = (46.3 : 42.7 : 9.4 : 1.6) | SM-102:Chol:(CDCL5:AP23):DSPC:DMG-PEG = (46.3 : 42.7 : (10.675 : 10.675) : 9.4 : 1.6) |
| **LNP Formula** | **C10 LNPs** | **AP23C5_25+ cLNPs** |
| lipid phase | Dlin-MC3-DMA:Chol:DOPE:DMG-PEG = (40 : 19.96 : 40 : 0.04) | SM-102:Chol:(CDCL5:AP23):DSPC:DMG-PEG = (40 : 19.96 : (4.99 : 4.99) : 40 : 0.04) |
